# Therapeutic validation of MMR-associated genetic modifiers in a human *ex vivo* model of Huntington’s disease

**DOI:** 10.1101/2023.12.05.570095

**Authors:** Ross Ferguson, Robert Goold, Lucy Coupland, Michael Flower, Sarah J Tabrizi

## Abstract

The pathological huntingtin (*HTT*) trinucleotide repeat underlying Huntington’s disease (HD) continues to expand throughout life. Repeat length correlates both with earlier age at onset (AaO) and faster progression, making slowing its expansion an attractive therapeutic approach. Genome-wide association studies have identified candidate variants associated with altered AaO and progression, with many found in DNA mismatch repair (MMR) associated genes.

We examine whether lowering expression of these genes affects the rate of somatic expansion in human *ex vivo* models using HD iPSCs and HD iPSC-derived striatal neurons. We have generated a stable CRISPR interference HD iPSC line in which we can specifically and efficiently lower gene expression from a donor carrying over 125 CAG repeats.

Lowering expression of each member of the MMR complexes MutS (MSH2, MSH3 & MSH6), MutL (MLH1, PMS1, PMS2 & MLH3) and LIG1 resulted in characteristic MMR deficiencies. Reduced MSH2, MSH3 and MLH1 slowed repeat expansion to the largest degree, while lowering either PMS1, PMS2 and MLH3 slowed it to a lesser degree. These effects were recapitulated in iPSC derived striatal cultures where MutL factor expression was lowered.

Here, reducing the expression of MMR factors by CRISPRi to levels typically reached by current therapeutics effectively slows the pathogenic expansion of the HTT CAG repeat tract. We highlight members of the MutL family as potential therapeutic targets to slow repeat expansion with the aim to delay onset and progression of HD, and potentially other repeat expansion disorders exhibiting somatic instability.

**GRAPHICAL ABSTRACT:** 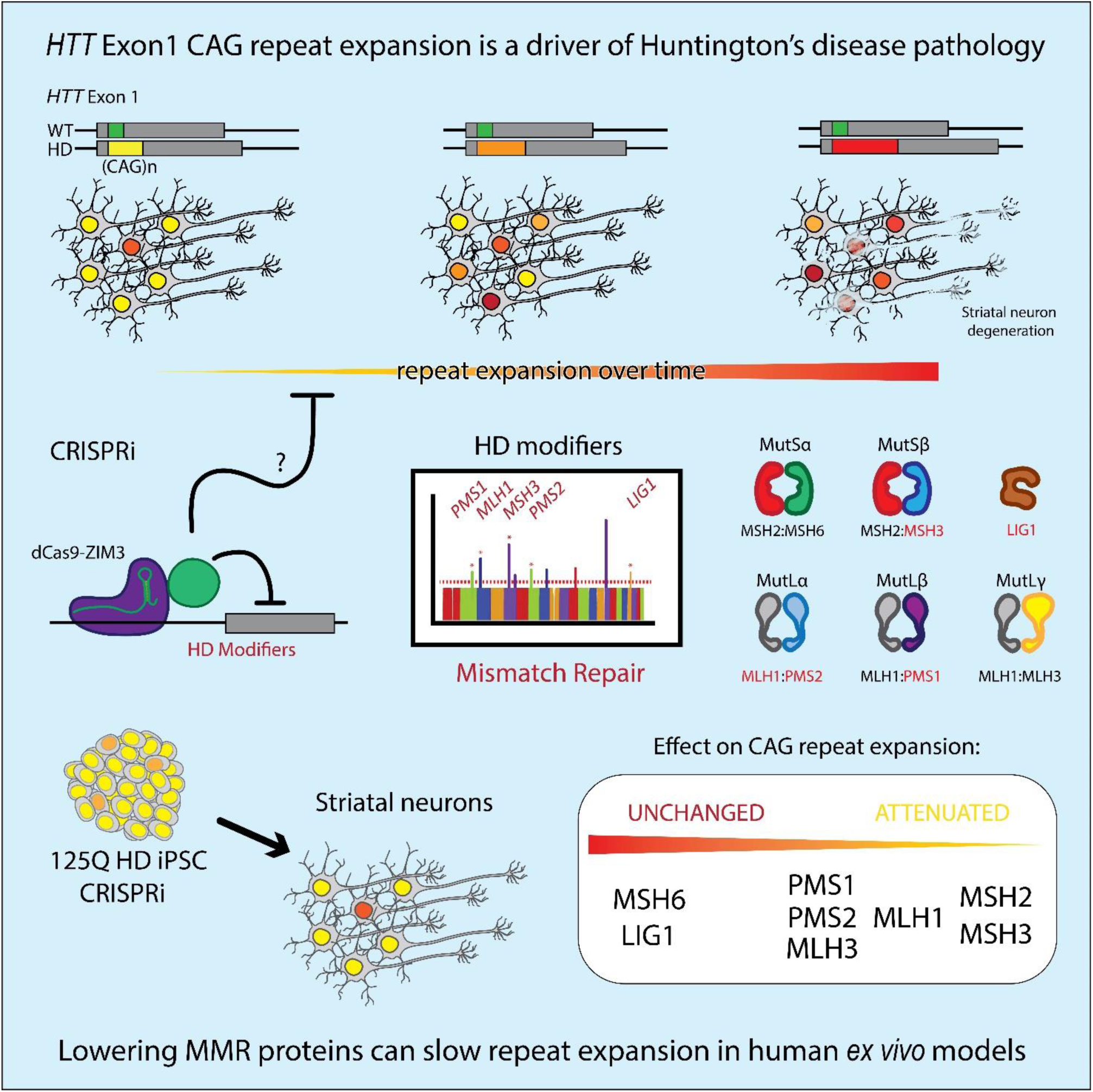

## INTRODUCTION

Huntington’s disease is a dominantly inherited fully penetrant neurodegenerative disease characterised by motor, cognitive and psychiatric symptoms with progression to death typically 15-20 years from onset. The mean age of onset is ∼43 years old and we currently have no disease modifying therapeutics. The onset and neurodegenerative pathophysiology of HD is driven by the inexorable increase in length of the expanded CAG repeat tract in exon 1 of the *huntingtin* (*HTT*) gene throughout the lifetime of the individuals carrying the pathogenic allele (Ciosi et al., 2019; Lee et al., 2019; MacDonald et al., 1993; Swami et al., 2009; Wexler et al., 2004). Here we examine candidate HD modifier genes identified through genetic studies, to validate targets to pursue in slowing repeat expansion rates, with the hope to delay disease onset and slow progression.

Expansion of the CAG repeat beyond a pathological threshold leads to progressive neurodegeneration (Donaldson et al., 2021). Striatal medium spiny neurons (MSNs) show particular vulnerability and degeneration is detectable well before onset (Albin, 1995; Mätlik et al., 2023; Morigaki and Goto, 2017; Pressl et al., 2023; Tabrizi et al., 2009; Vonsattel et al., 1985). Striatal MSNs and layer V cortical pyramidal cells more frequently carry high repeat length burdens than other neuron sub-types in both mouse and human (Gonitel et al., 2008; Kennedy et al., 2003; Mätlik et al., 2023; Pressl et al., 2023), suggesting increased repeat expansion is a significant driver of vulnerability in these neurons. Although CAG repeat length correlates with its expansion rate, additional cell-type dependent modifiers that drive somatic CAG repeat mosaicism within an individual are not understood (Kacher et al., 2021; Kennedy et al., 2003; Shelbourne et al., 2007).

Increasing length of the *HTT* CAG repeat robustly correlates with earlier age at onset (AaO) and disease progression, however substantial variation exists between HD carriers with similar repeat lengths (Ciosi et al., 2019; GeM-HD Consortium, 2019; Wright et al., 2019).. Genome wide association studies (GWAS) have identified variants associated with altered AaO and progression in HD cohorts controlled for repeat length (Bettencourt et al., 2016; Ciosi et al., 2019; GeM-HD Consortium, 2015, 2019; Moss et al., 2017). Many variants are located in DNA damage response (DDR) genes. Variants in the Fanconi-anaemia pathway associated protein FAN1 strongly associated with altered AaO, and FAN1 has been convincingly shown to significantly alter repeat expansion in multiple models (Goold et al., 2019, 2021; Loupe et al., 2020). Disease modifying variants in multiple members of the mismatch repair (MMR) pathway were also identified, including MSH3, MLH1, PMS1, PMS2 & LIG1.

Mismatch repair has long been implicated in instability at the *HTT* CAG repeat (Manley et al., 1999). Effective MMR in mammalian cells relies on the MutS heterodimer complexes — either MSH2 and MSH6 (MutSα) or MSH2 and MSH3 (MutSβ). While MutSα primarily engages 1-2 base mismatches, MutSβ recognizes larger extra-helical extrusions, though with some overlap (Genschel et al., 1998; Iyer et al., 2006). *In vitro* studies propose that MutSβ associates with extra-helical DNA structures similar to those which occur at highly repetitive sequences like the *HTT* CAG repeat (Lang et al., 2011). Engagement of a lesion by either MutS complex recruits MutL protein heterodimers to form ternary complexes. MLH1 serves as the common component in each MutL heterodimers, partnering with either PMS2 (MutLα), PMS1 (MutLβ), or MLH3 (MutLγ) (Iyer and Pluciennik, 2021). Upon activation by MutS and PCNA, MutLα incises DNA flanking the mismatch (Genschel et al., 2017), while MutLγ nicks in the opposing strand (Kadyrova et al., 2020). Unlike PMS2 and MLH3, PMS1 lacks a nuclease domain so MutLβ function in MMR is unknown. At conventional MMR substrates exonuclease EXO1 resects the lesion-containing strand, which then undergoes resynthesis and ligation facilitated by DNA polymerase δ and LIG1 (Iyer and Pluciennik, 2021).

Substrate, kinetics, ternary complex formation, and the co-factors present at unstable repeats are not well understood in comparison to non-repeat MMR lesions, yet additional repeats are inevitably incorporated. Loss of individual MMR proteins in mouse models significantly affects repeat expansion: Loss of Msh3 arrests repeat expansion in HD and myotonic dystrophy mouse and cell models (Goold et al., 2021; Keogh et al., 2017; Lee et al., 2010; Manley et al., 1999; Owen et al., 2005; Tomé et al., 2009; Wheeler et al., 2003), while Msh6 loss has minor variable effects (van den Broek et al., 2002; Du et al., 2012). Likewise, loss of Mlh1, Pms1 and Mlh3 slows repeat expansion in HD and Friedrich’s ataxia (FRDA) models (Miller et al., 2020; Pinto et al., 2013) while Pms2 both promotes and supresses expansion in different models (Bourn et al., 2012; Gomes-Pereira et al., 2004; Halabi et al., 2018; Miller et al., 2020).

The process of somatic repeat expansion in the brain appears to be a major factor in driving disease progression, leading to production of toxic HTT protein species. Hong et al., (2021) suggest that rather than a continuum of repeat length dependent toxicity there is a critical threshold beyond which severe dysfunction occurs in vulnerable cells. Slowing the rate of expansion is clearly an attractive target for therapeutic intervention in HD, and other repeat expansion disorders which exhibit somatic instability, but first it is necessary to confirm which disease-modifying candidates actually play a role in the expansion process. The occurrence of gene-function modifying variants within the population, as identified by GWAS, suggests their effects are tolerated to some degree. Therefore, targeted expression modulation of these genes could also likely be tolerated in individuals. For example, functionally deleterious polymorphisms in *MSH3* slow repeat expansion and HD progression (Flower et al., 2019), and lowering MSH3 expression is currently being pursued therapeutically (O’Reilly et al., 2023).

Here, we have asked whether lowering expression of candidate MMR genes to levels achievable by current therapeutics can affect the rate of repeat expansion in a human *ex vivo* model of HD. We investigated HD modifier MMR genes and their cognate heterodimer partners by targeting MutS (MSH2, MSH3 & MSH6), MutL (MLH1, PMS1, PMS2 & MLH3), and LIG1 expression using a CRISPR interference system in HD iPSCs. This lowers expression by targeting a catalytically inactive Cas9 fused to a transcriptional repressor to the transcriptional start site of each gene. The effects of lowering each target on repeat expansion were then assayed both in dividing HD iPSCs and HD iPSC derived post-mitotic striatal neurons.

## MATERIALS & METHODS

### Cell culture

HD 125Q iPSCs (Goold et al., 2021) were maintained in E8 Flex (Gibco) on Geltrex (1:100, Gibco) coated plasticware (Nunc) and passaged using 0.5mM EDTA in PBS (Gibco). U2OS cells were maintained in DMEM supplemented with 10% FBS, 1% L-glutamine and 1% pen/strep (Gibco) and passaged using Trypsin/EDTA.

### U2OS repeat expansion assay

U2OS cells were transduced with p’HRsincpptUCOE HTT Ex1 118Q IRES eGFP, and cultured for the indicated time periods and changes in CAG repeat length determined by fragment analysis, as previously reported (Goold et al., 2019).

### Striatal differentiation of iPSCs

Differentiation was performed based on Arber et al., 2015. iPSCs were passaged 1:2 onto Matrigel coated plates to become confluent within 24-48h with feeding every 24h. Once confluent, cells were washed and switched to neural induction media (DMEM:F12 & Neurobasal A 1:1 (both Gibco) with 1x L-glutamine (Gibco), 1x PenStrep (Gibco), 0.5x N2 (Gibco), 0.5x B27 without vitamin A (Gibco), 100μM β-mercaptoethanol with 100nM LDN-193189 (Sigma) and 10μM SB431542 (Cambridge Biosciences)). Media was changed every 24h up to day 10 on which cells were mechanically clump passaged using 0.02% EDTA in PBS onto fibronectin coated plates and maintained in the same media without SMAD inhibitors and including 25ng/ml activin-A (Peprotech). All subsequent media changes were half volumes and every 48h before passaging again onto poly-l-ornithine and laminin coated 12w plates on day 20. After day 26, B27 was switched to include vitamin A and both BDNF and GDNF included at 10ng/ml (Peprotech). Activin-A was omitted from day 36 with half media changes every 72h.

### Design, cloning and preparation of CRISPRi guides

CRISPRi sgRNAs for knocking down MutL component expression were designed and compared using the online tools CRISPick (portals.broadinstitute.org/gppx/crispick/) and CRISP-ERA (crispr-era.stanford.edu/) (Table S1). Non-targeting control guide sequences were obtained from Sanson et al., (2018). Corresponding oligo pairs were hybridized and inserted into the pLentiGuide vectors by GoldenGate cloning using BsmBI-V2 and T4 Ligase (NEB) then transformed in NEB stable cells. Isolated clones were screened by colony PCR across the stuffer insert and confirmed by restriction digest followed by sequencing from the U6 promoter. Primer sequences can be found in Table S1. Endotoxin free plasmid for transfection was purified using PureYield column kits (Mini/Maxi, Promega). Plasmid DNA for direct use in electroporation of iPSCs was further concentrated by precipitation. Plasmid DNA used to generate lentiviral particles were transfected into HEK293 cells using LentiX single-shot VSV-G (Takara) following manufacturer’s instructions. Harvested virus was concentrated by centrifugation through a 10% sucrose cushion at 10,000g for 4h (Jiang et al., 2015). Viral titre was determined by qPCR (LentiX, Takara).

### Modification of guide and dCas9-KRAB constructs

The KOX1 KRAB domain of the dCas9-KRAB fusion in the CLYBL targeting vector pC13 dCas9-TagBFP-KRAB (Tian et al., 2019, addgene #127968) was swapped for ZIM3 KRAB domain from pLX303-ZIM3-KRAB-dCas9 (Alerasool et al., 2020, addgene #331490). The ZIM3 coding sequence was amplified by PCR using as pLX303 as template. A 5’ HindIII site and a 3’ FseI site were appended by inclusion in the primer sequences. The KRAB domain in the dCas9-tagBFP-KRAB degron construct pRT029 was dropped out by HindIII/FseI digest and replaced by the HindIII/FseI digested ZIM3 amplicon. ZIM3 and TagBFP were amplified with overlapping homology with the pC13 5’ BamHI and 3’ MluI sites. The final pC13 dCas9-BFP-Zim3 construct was assembled from these fragments using HiFi assembly mix (NEB).

pLentiGuide-Hygro-eGFP (Ho et al., 2011, addgene #99375) was modified to include PiggyBAC inverted terminal repeats flanking the sgRNA and HygroR-eGFP expression cassette. A backbone fragment containing AmpR was excised by FspI. The same fragment was amplified by PCR using primers appended with the PB TIR sequences and blunt cloned back into the same sites.

All restriction enzymes were purchased from NEB. All PCR based cloning steps were performed using KAPA HiFi taq (Roche) and verified by sequencing, primer sequences can be found in Table S2. An expression vector for the hyperactive PiggyBAC transposase with an mCherry reporter (pRP EF1A-hyPBase CMV-mCherry) was synthesised by Vectorbuilder.

### TALEN-mediated knock-in of dCas9-ZIM3 in 125Q HD iPSC

The modified pC13 dCas9-BFP-ZIM3 construct with CLYBL homology arms was co-nucleofected into 125Q iPSC with two additional constructs carrying TALENs targeting the CLYBL locus (pZT-C13-R1 and pZT-C13-L1, addgene #62196, #62197 (Cerbini et al., 2015)) in P3 solution using program CA137 in a 4D-Nucleofector (Lonza). Cells were seeded at clonal density and selection in G418 was initiated 48 hours later. Colonies were picked and screened for BFP expression. PCRs were performed spanning both the 3’ and 5’ homology arm junctions to identify integration at the intended sites.

Zygosity was determined by qPCR using genomic DNA as template using primers for Cas9 and ZNF80. A two-copy reference was created by cloning the ZNF80 amplicon into the AleI site in pLX303. Copy number qPCR was performed using Forget-Me-Not qPCR Master Mix (Biotium) on a QuantStudio5 system.

### gRNA introduction by lentiviral transduction

iPSCs were transduced at 70% confluence with ∼50 MOI in the presence of 10µg/ml polybrene in 24 well plates. After 24-48h cells were passaged to 6 well plates and selection initiated in 50µg/ml hygromycin. Selection was maintained for two weeks before assaying knock-down.

### gRNA introduction by PiggyBAC transposition

iPSCs at 70% confluence were preincubated for 30min with ROCKi (10µM Y-27632, Sigma) then dissociated to single cells using TrypLE (Gibco). Cells were resuspended in complete media with ROCKi, centrifuged at 250g for 3min then resuspended in PBS and counted. 2x10^5^ cells/ml were nucleofected in 20µl using nucleocuvette strips with 0.5µg guide plasmid, 0.5µg hyPBase plasmid and 0.1µg BCL-XL plasmid (Li et al., 2018). After 24-48h cells were passaged to 6 well plates and selection initiated in 50µg/ml hygromycin. Selection was maintained for two weeks before assaying knock-down.

### Generation of CRISPR/Cas9 knock-out cell lines

MSH3 and MLH1 were knocked-out in U2OS cells previously (Goold et al., 2021). MSH6 knock-out U2OS and 125Q iPSCs were generated by nucleofection with complexed HiFi Cas9 and paired Alt-R guide RNAs (IDT) targeting exon 2 or exon 4 (Table S3). For both methodologies, single cell clones were isolated, expanded and screened by PCR for the deletion of the guide flanked sequence (Table S2) then knockout was confirmed by Sanger sequencing and Western blot.

### RT-qPCR

Total RNA was prepared by Qiagen RNAeasy columns from cells lysed directly in wells in RLT buffer. cDNA synthesis performed using the SuperScript™ IV First-Strand Synthesis System with 1:1 Oligo dT and random hexamer primers (all Thermo). qPCRs were set up using 10ng template in 15ul reactions with TaqMan Fast Advanced mastermix (Thermo) with assays in two-plex (Table S4) on a Quantstudio 5. All qPCRs were performed in technical triplicate using exon spanning assays and presented as fold change relative to the geometric mean of three references genes (UBC, ATP5B & EIF4A2).

### Western Blot

Frozen cell pellets were lysed in two pellet volumes of RIPA (Sigma) with protease inhibitors (Halt, Promega) by trituration followed by a twenty minute incubation on ice with Benzonase. (Sigma). Protein concentration was determined by BCA assay (Pierce) and 20µg of protein were run on 4-12% Bis-Tris NuPage gels (Thermo) against Spectra Multicolour high-range markers (Thermo) before transfer to 0.2µm nitrocellulose overnight using an XCell ii system at 35V (Thermo). Membranes were blocked in 5% milk in PBS for one hour before incubating with primary antibodies overnight at 4°C with shaking. After four washes of ten minutes each with PBST, membranes were incubated with secondary antibodies for one hour at room temperature. After four further washes with PBST and two with PBS, membranes were imaged on a Licor Odyssey CLx. Band intensity of proteins of interest are presented relative to ACTB and normalised to non-targeting controls. Antibodies and dilutions can be found in Table S5.

### Immunocytochemistry

Cells cultures in 96 well PhenoPlates (PerkinElmer) were fixed in 4% PFA in PBS for fifteen minutes at RT, followed by three PBS washes of ten minutes each. Fixed cells were incubated in blocking buffer comprised of 5% BSA (Sigma), 1% FBS (Gibco) and 0.5% Triton-X100 (BDH) in PBS. Samples were incubated with primary antibodies overnight in diluent (blocking buffer 1:10 with PBS). After four washes of ten minutes with PBST, samples were incubated with secondary antibodies and Hoechst counterstain for two hours RT. Following four further ten minutes PBST washes and two PBS washes samples were prepared for microscopy with 90% glycerol and 0.5% N-propyl gallate in 0.1M Tris pH 7.4. Images were acquired on an Opera Phenix (Perkin Elmer) from at least four random fields in three wells from each experiment. Images for quantification were acquired with identical setting with flat field correction and further analysed using Fiji (Schindelin et al., 2012). Normality was determined using the D’Agostino-Pearson test and tested by ANOVA and Bonferroni post-hoc in Prism 9 (GraphPad).

### Fragment analysis

DNA was prepared by either QuickExtract (Lucigen) from iPSCs or Qiagen DNeasy columns (MSNs and U2OS) following manufacturer’s instructions. PCR across the *HTT* CAG repeat was performed on 10ng gDNA using AmpliTaq Gold 360 (Thermo) with 6-FAM labelled HD3F CCTTCGAGTCCCTCAAGTCCTT (Teo et al., 2008) and HD5 Mangiarini CGGCTGAGGCAGCAGCGGCTGT (Mangiarini et al., 1996), with the following cycling conditions: 94°C for 90 s, 35 cycles of 94°C for 30 s, 65°C for 30 s and 72°C for 90 s, and the final extension at 72°C for 10 mins. 2µl of PCR product was denatured in 10µl HiDi formamide (Thermo) with 0.5µl Mapmarker ROX 1000 (Eurogentec) and run on an ABI 3730XL genetic analyser. Sequencer output was processed using GeneMapper (Thermo) and a custom R script available at https://michaelflower.org using a threshold of 20% of modal peak height and a correction factor of 2.724117. Data presented as modal repeat length relative to control samples and a modified instability index (Lee et al., 2010), calculated here as the sum of the change in repeat length relative to the mode, multiplied by proportional peak height, for each peak in the distribution but measured relative to the control repeat length. Data was analysed by linear mixed-effects model with post-hoc comparison by Tukey’s method in Prism 9 (GraphPad).

### Genetic constraint in selected HD modifier and DNA repair genes

To measure genetic constraint, or the gene’s intolerance to variation, we analysed loss of function (pLoF) metrics from the gnomAD database (v4.0.0, 807,162 samples) (Gudmundsson et al., 2022). This analysis calculated the ratio of observed / expected (o/e) number of loss-of-function variants, using transcripts annotated by GENCODE v39 and the GRCh38 reference. The expected counts are based on a model that accounts for sequence context and methylation. pLoF values are presented along with the 90% confidence interval. When a gene has a low pLoF value, it is under stronger selection. For example, a pLoF value of 0.2 means that there were 20% of the expected number of variants observed in that gene. Whilst pLoF values are continuous, scores below 0.6 are generally accepted to indicate significant selection against loss-of-function variation.

## RESULTS

### Construction of a stable CRISPRi HD iPSC line

We constructed a CRISPRi HD iPSC line in which we could both effectively lower target gene expression and measure somatic instability at the *HTT* CAG repeat. A dCas9-KRAB repressor construct was stably integrated into the CLYBL safe-harbour in chromosome 13 of our highly characterized 125Q juvenile HD iPSC line (Goold et al., 2021). We modified the CLYBL targeting construct of Tian et al., (2019) to swap the KOX1 KRAB domain for the more effective ZIM3 KRAB domain identified by Alerasool et al., (2020). (Figure 1A). We compared these fusions using *PMS2* targeting guides and found ZIM3 mediated repression was equal to or greater than that observed with the same guide sequences paired with the KOX1 KRAB fusion (Figure S1A).

**Figure 1.**
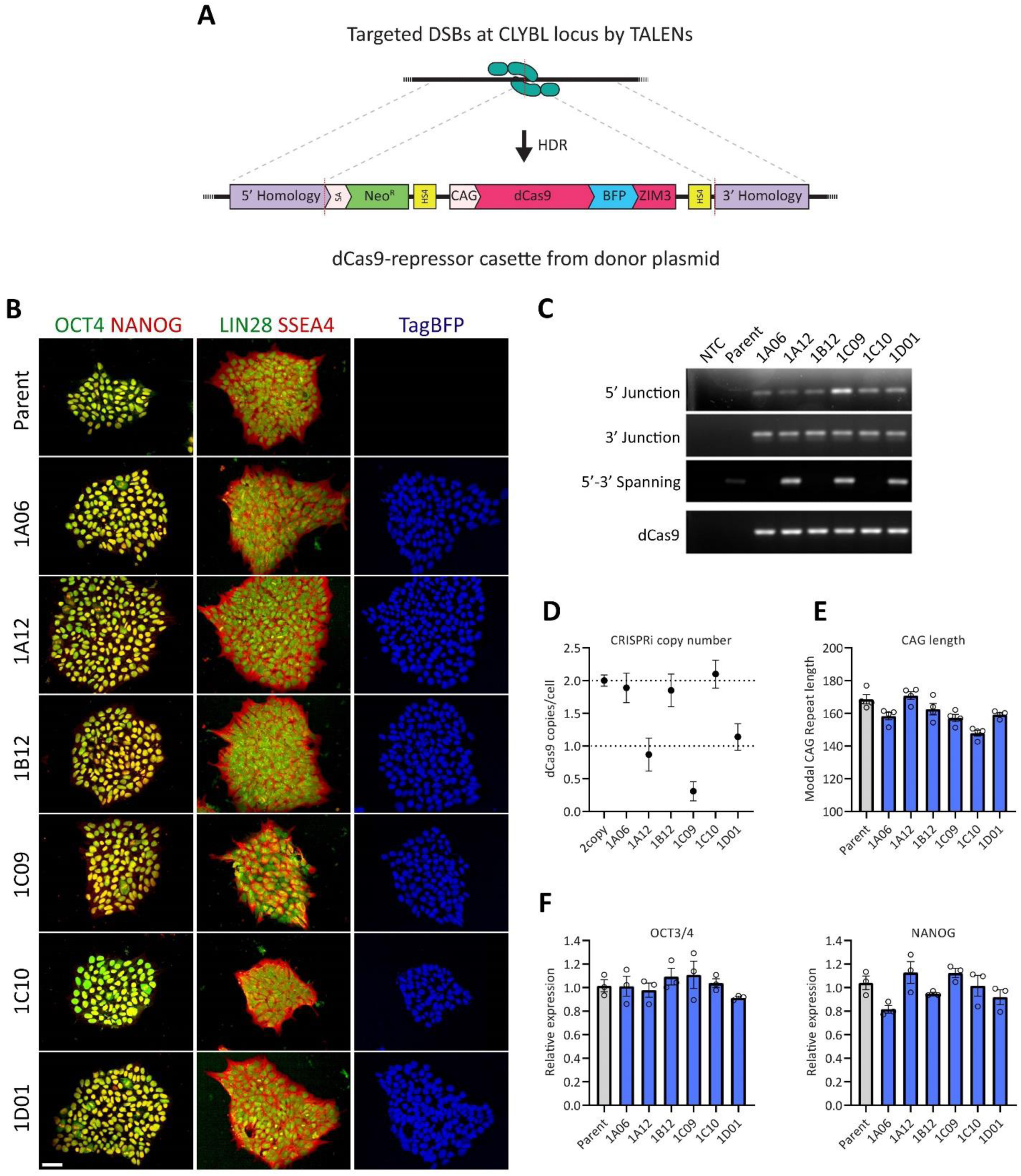
Construction and validation of the 125Q HD CRISPRi iPSC lines. Schematic showing TALEN-mediated knock-in of a dCas9-tagBFP ZIM3 KRAB construct into the CLYBL safe-harbour (A). Single cell clones were isolated and screened for TagBFP expression and pluripotency associated marker expression. Scale bars 50µm (B). Accurate knock-in was verified by PCR across homology arm junctions (5’ and 3’ junctions) and insert site spanning (5’ to 3’ spanning) PCR alongside an internal dCas9 PCR (C). Zygosity and off-target integration was assessed by copy number qPCR (D). Modal CAG length was determined by fragment analysis (E). Expression of pluripotency associated genes was further assessed by qPCR for OCT3/4 and NANOG relative to the parental iPSC line (F). Mean values ±SEM in triplicate.

Targeted integration of the construct at the CLYBL locus was directed by TALEN mediated double strand breaks and subsequent homology directed repair using the homology arms in the dCas9-repressor donor construct (Figure 1A). TagBFP expression was used to screen single cell iPSC clones for successful knock-in after selection in G418 (Figure 1B). We further screened six of these clones by PCR to amplify an internal Cas9 product, a product spanning the 5’ and 3’ junction between the CLYBL locus and the donor construct, and a product spanning the TALEN targeted cut site (5’-3’ spanning, Figure 1C). These PCRs confirmed knock-in at the correct site in all clones and indicated three of the six clones were homozygous for the CRISPRi construct. This was further confirmed by copy number qPCR (Figure 1D) which also highlighted one of the three potential heterozygous clones as potentially mixed / non-clonal. The range of modal CAG length of the knock-in clones remained within ∼20% parent population. (Figure 1E). Clones maintained good iPSC-like morphology, growth, and pluripotency-associated marker expression (Figure 1BF). *OCT3/4* and *NANOG* transcript levels were not significantly different to those of the parent 125Q line by qRT-PCR (Figure 1F). Immunostaining for the pluripotency markers OCT4, NANOG, LIN28 and SSEA4 showed expected distributions and levels comparable to that of the parent line (Figure 1B). TagBFP distribution showed the dCas9-TagBFP-ZIM3 fusion protein was localised to the nucleus as expected (Figure 1B).

### Identification of guide sequences which effectively lower MMR factor expression

Multiple guide sequences were selected that target the transcriptional start site of each gene of interest: *MSH2*, *MSH3* & *MSH6* (MutS components), *MLH1*, *PMS1*, *PMS2* & *MLH3* (MutL components), as well as *LIG1* (Figure 2A). Oligos carrying these guide sequences were cloned into pLentiGuide based expression constructs. (Figure S1B). These constructs also carry EGFP and hygromycin resistance markers and were introduced to the CRISPRi HD iPSC line either by lentiviral transduction (MutL) or PiggyBAC transposition (MutS & LIG1) (Figure 2B). Both methods resulted in comparable initial efficiencies by EGFP expression with ∼20-30% GFP^+^ cells at 24h post-introduction.

**Figure 2.**
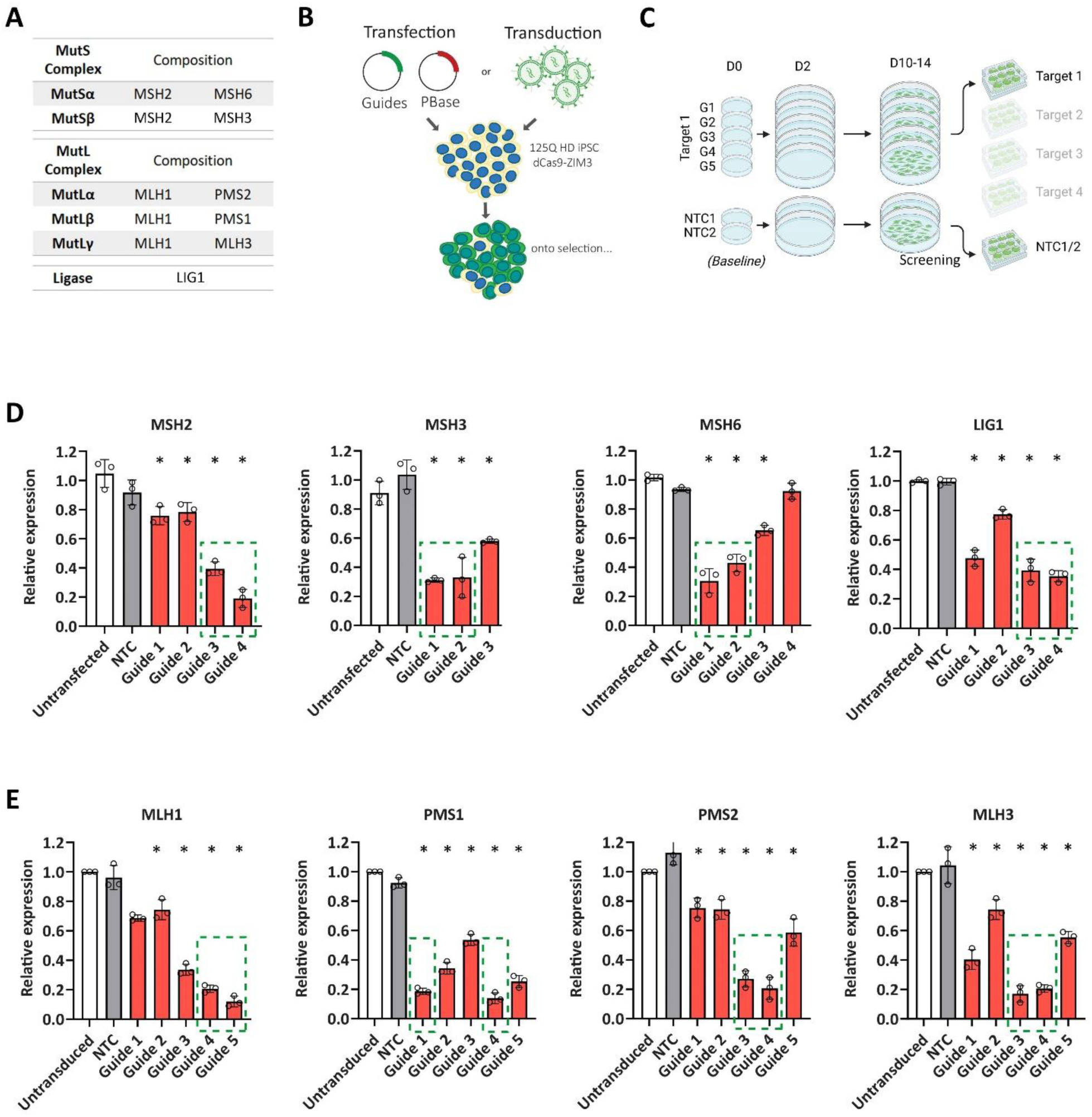
Effective knockdown of MMR factors after introduction of sgRNAs to a 125Q HD CRISPRi line. Guide sequences were introduced into the dCas9-repressor iPSC line either through lentiviral transduction or co-electroporation with the PiggyBAC transposase, followed by selection (A). Guides targeting the MutL factors MLH1, PMS1, PMS2 and MLH3 were introduced by lentivirus, while MutS factors MSH2, MSH3 & MSH6 as well as LIG1 were introduced by electroporation and PiggyBAC transposition (B). Up to five guides per target and two non-targeting (NTC) guides were used in each round and their efficacy in lowering target expression was assayed after selection in hygromycin (C). Transcript levels were determined by qPCR for MutS & LIG1 (D), and MutL (E) normalised to un-transfected/transduced controls. Data compared by ANOVA with Bonferroni’s post hoc. Error bars show SEM. N=3, error bars ±SEM, *= P<0.05. Two guides per target were taken forward, highlighted by green dashed boxes.

Up to five guides per target were introduced in parallel alongside non-targeting guides (NTC1&2) (Figure 2C). After selection and expansion in hygromycin containing medium, target expression was assayed at the transcript level by qPCR. Significantly lower target transcript levels were seen with each guide in comparison to the conditions without guides or with non-targeting guides. Lowering of transcript levels to around 70% was seen for the top MSH2 and MSH3 guides while MSH6 and LIG1 guides achieved around 60-70% lowering (Figure 2D). At least two guides for each MutL component (MLH1, PMS1, PMS2 or MLH3) lowered transcript levels by around 80% (Figure 2E). From these sets, two guides with good efficacy were selected to characterise the consequences of lowering each target on CAG repeat expansion (Figure 2DE, dashed boxes). First however, they were characterised in terms of effect on target protein levels and MMR proficiency.

### CRISPRi mediated lowering of MutS & MutL factors in iPSCs leads to characteristic MMR deficiencies

Homogeneous expression of EGFP from the guide construct can be seen across the non-targeting and targeting guides after selection (Figure 3A). Immunocytochemistry for each CRISPRi MMR protein target shows a primarily nuclear distribution in iPSCs, with some variation in intensity likely linked to cell cycle status (Schroering et al., 2007). This pattern remains where target transcript levels are lowered by CRISPRi, though at substantially reduced intensity (Figure 3A).

**Figure 3.**
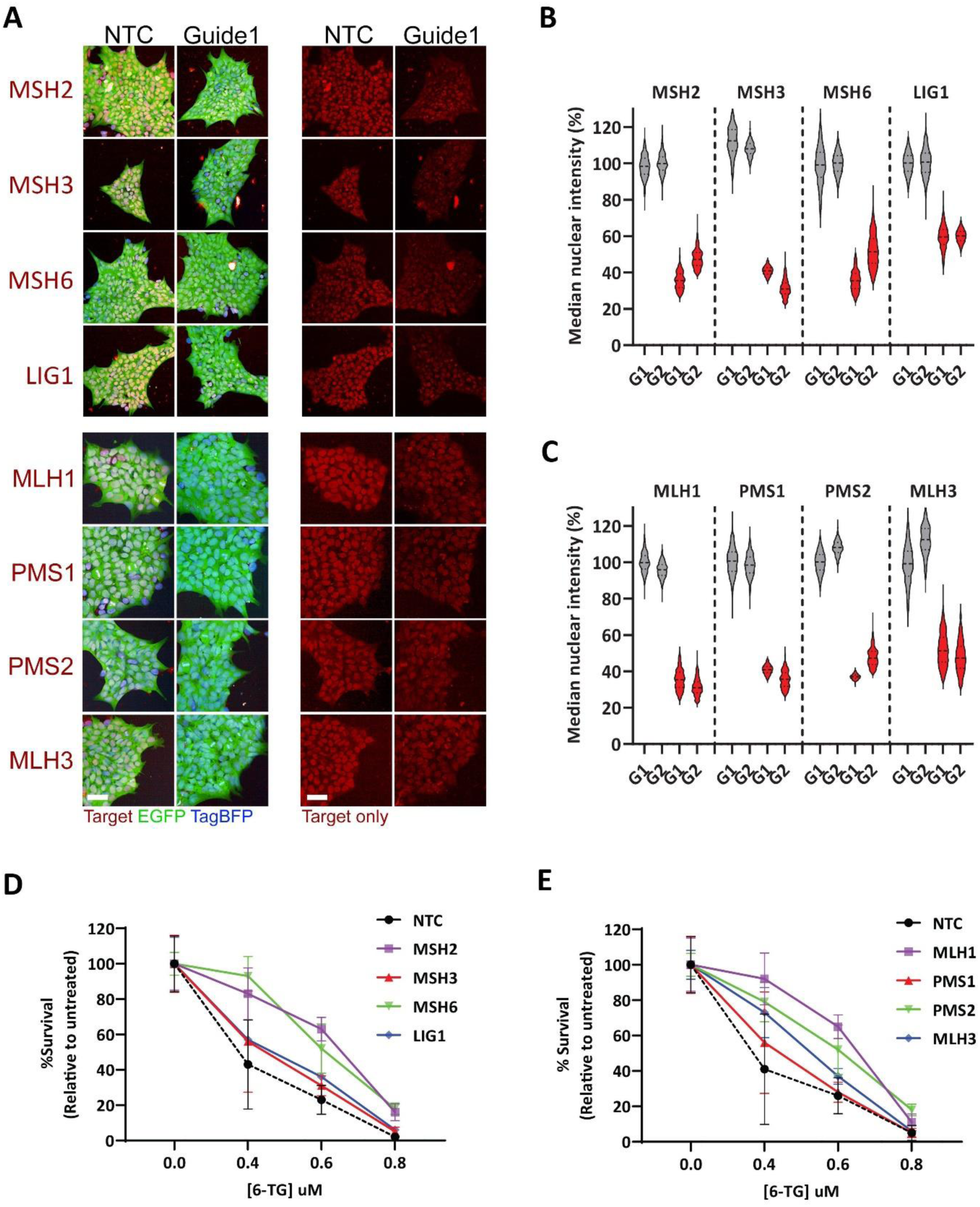
Homogeneous lowering of MMR factor expression in CRISPRi pools leads to characteristic MMR deficiencies. Immunostaining for indicated target proteins in red channel shows knockdown is consistent cell-to-cell (A). Representative images for a non-targeting control guide and a targeting guide. EGFP from the guide vectors and TagBFP from the dCas9-KRAB knock-in. Scale bars 50µm. Quantification of fluorescence intensity for each target normalised to NTC1 levels for MutS & LIG1 (B), and MutL (C). Line indicated median values, 1-2000 cells per plot from four random positions in three independent experiments. iPSC colony formation after treatment with the indicated concentration of 6-thioguanine (6-TG) for CRISPRi lowered MutS & LIG1 (B), and MutL (C). Mean colony frequency ±SEM in triplicate as a proportion of untreated cells of the same genotype, normalised to NTC.

Quantification of the median nuclear intensity of each target within MutS & LIG1 (Figure 3B) and MutL (Figure 3C) confirms the homogeneity of the knock-down and correlates with the reduction in transcript level seen for each guide (Figure 2EF, dashed boxes). Significant and consistent lowering is seen between cells within each knock-down population.

Total loss, reduced levels or functional mutations in the various members of the MMR pathway lead to characteristic defects in MMR response (Li, 2008). We have assayed MMR proficiency here using survival after treatment with the base analog 6-thioguanine (6TG). The incorporation of 6TG during DNA synthesis leads to MMR-dependent cell-cycle arrest, futile repair attempts and cell death whereas in the absence of MMR cells survive (Yan et al., 2003). Non-targeting control iPSCs and those with CRISPRi lowered MMR factor expression were seeded at clonal density and treated with a range of 6TG concentrations for 24h followed by quantification of the frequency of colonies formed after one week.

The small single base lesions associated with 6TG incorporation would be repaired primarily through MutSα (MSH2:MSH6) dependent means. As expected, the loss of the core MutS and MutL factors had the largest effect on survival, as did lowering MSH6 (Figure 3DE). In contrast, lowering MSH3 or LIG1 had no significant effect on 6TG sensitivity in comparison to controls. Lowering the MutL co-factors PMS2 and MLH3 had an intermediate effect in comparison to lowering their partner MLH1 while lowering PMS1 showed only a very minor difference in survival to controls (Figure 3DE).

CRISPR-Cas9 mediated knock-out of MSH6, MSH3 and MLH1 in U2OS osteosarcoma cells resulted in the complete loss of each protein (Figure S2A and Goold et al., (2021)). Both MSH6 and MLH1 null U2OS cells showed improved survival post-6TG treatment, in contrast to wild-type and MSH3 null cells (Figure S2B). Similarly, the same MSH6 knock-outs made in the 125Q iPSC line (Figure S2C) resulted in the most robust survival post-6TG treatment, indicating that the effect seen upon lowering by CRISPRi in iPSCs likely does not represent the maximal MMR deficiency possible in each case (Figure S2D).

The lack of effect seen where LIG1 has been lowered may suggest either further lowering is required or there is redundancy with from LIG family members, such as LIG4 whose levels remain unaffected (Figure S3). Having shown that lowering MutS or MutL levels to the degree achieved here is sufficient to detect an MMR phenotype, we went on to characterise their effect on somatic expansion in dividing iPSCs and post-mitotic MSNs.

### Lowering MutSα and each MutL component slows somatic expansion in dividing iPSCs

We measured the effect of target lowering on somatic instability in iPSCs over the course of eighty days during routine passaging. CAG repeat length was measured by fragment analysis at twenty-day intervals and changes in modal CAG length or instability index (Lee et al., 2010) were reported relative to the baseline at the start of the time course (Figure 4A). No substantial changes in target transcript levels were observed across the time course by qPCR, indicating no significant silencing of any CRISPRi component was occurring (Figure 4B).

**Figure 4.**
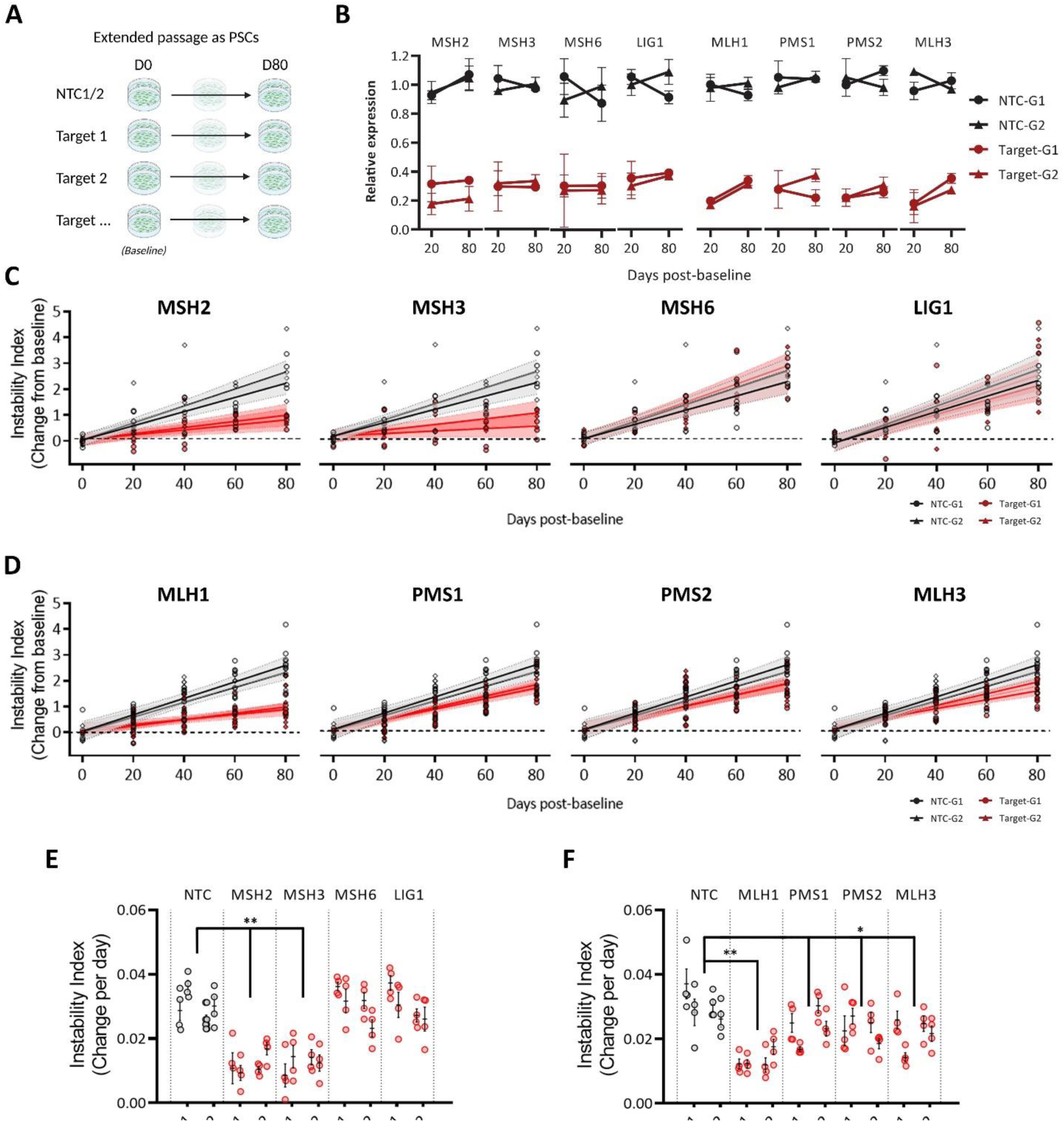
Reduced expression of MutS and MutL components slows somatic expansion in dividing iPSCs. Somatic instability at the pathogenic CAG repeat tract in the 125Q HD iPSC line can be measured over time in across routinely passaged cultures, relative to the starting point at day 0 (D0, baseline) (A). qPCR for each target shows lowering persists over the time-course (B). Mean expression relative to day zero ±SEM in triplicate for two NTC guides and the selected two target specific guides at day 20 and 80. Instability index relative to control over an eighty-day period for MutS & LIG1 (C), and MutL (D) lowered cultures relative to baseline on D0. Four cultures per guide passaged in parallel each from two independent CRISPRi pools, dashed lines 95% confidence intervals. Rate of change in instability index relative to control per day for MutS & LIG1 (E), and MutL (F). Open circles represent parallel cultures with mean bar ±SEM. * P<0.05 * P<0.005.

Over the eighty-day time-course we saw an increase in modal CAG length of 4-6 from baseline (Figure S4A) and an increase in instability index by around two units in NTC guide cultures (Figure 4C). Across the same period, an increase of less than one unit was measured in the instability index of cultures where expression of either MSH2 or MSH3 was lowered (Figure 4C, P<0.005), accompanied by an modal increase of only 1-2 CAG units (P<0.0005, Figure S4A). In contrast, no significant change in modal CAG repeat length or instability index was found between control cultures and those with lowered MSH6 or LIG1 (Figures 4C and S4A). Where MutL expression was targeted (Figure 4D) we observed the slowest rates of expansion where MLH1 levels were lowered, with a change in instability index relative to baseline around one unit (P<0.005) and an increase of only ∼1 CAG by day 80 (P<0.005, Figure S4A). Lowering PMS1, PMS2 and MLH3 also had a small but significant effect, reducing instability index by around half a unit and modal repeat length by ∼1-2 CAGs over the full time-course in comparison to control cultures (P<0.05, Figure 4D and S4A). Rates of change in instability index and modal CAG per day were derived from the mixed models of the four independently passaged cultures per guide and transduction. Lowering MSH2, MSH3 and MLH1 reduced the rate of expansion in iPSCs by between 60 and 65% (Figure 4EF) while lowering PMS1, PMS2 or MLH3 reduced the rate by between 25 and 35% compared to control rates (Figure 4F).

These results matched data obtained using a second model of repeat expansion in U2OS cells. In these cells, repeat expansion can be measured over time in a 118Q repeat present in a *HTT* exon1 expression construct delivered by transduction (Goold et al., 2019, 2021). The effect on repeat expansion in this construct was measured in MSH3, MSH6 and MLH1 knock-out cells over a 40-day period (Figure S5). As expected, the total absence of either MSH3 or MLH1 resulted in an almost complete arrest of expansion while loss of MSH6 had no significant effect. Rescuing expression using an inducible strep-tagged variant of either MSH3 or MLH1 restored repeat expansion to wild-type rates (Figure S5).

### Lowering of each MutL component slows somatic expansion in striatal neuron cultures

We next differentiated a subset of CRISPRi iPSCs to striatal neuron identity to investigate the effect on somatic expansion in populations of post-mitotic neurons. These cultures are enriched for medium spiny neurons (MSNs), known to be vulnerable to repeat expansion during disease (Swami et al., 2009; Telenius et al., 1994). We did not pursue the effects of lowering MSH6 or LIG1 in neurons as neither showed a significant effect in iPSCs using this system. Although lowering MSH2 or MSH3 resulted in a robust slowing of repeat expansion in iPSCs, lowering of MSH2 *in vivo* is unlikely to be a viable long-term approach due its oncogenic potential (Dominguez-Valentin et al., 2020; Therkildsen et al., 2015). In contrast, loss of MSH3 is substantially less oncogenic, particularly in the context of CNS cancers and multiple approaches to therapeutic MSH3 lowering are currently under investigation (Flower et al., 2019; O’Reilly et al., 2023). We instead chose to focus on characterising the effect of lowering the less well understood MutL factors – MLH1, PMS1, PMS2 and MLH3. Comparison of the expected frequency to actual occurrence of variants in MutL factors that are predicted to impair function in the population suggests variants in PMS1 and PMS2 to be well tolerated, as well as MLH3 and MLH1 to a lesser degree (Figure S6).

Constitutive lowering of each MutL factor does not compromise the capacity of the iPSCs to differentiate into striatal neurons. Immunocytochemistry for key markers of neuronal and specifically striatal identity neurons showed robust and comparable frequencies between cultures of each set of lowered factors at day 100 (Figure 5A). Quantification of the post-mitotic neuron marker NeuN showed a consistent 70-90% frequency between neuronal cultures of each genotype (Figure 5B). Similarly, quantification of the MSN marker DARRP-32 ranged between 30-40% of the total population across each genotype (Figure 5C), as expected for the differentiation methods used (Arber et al., 2015).

**Figure 5.**
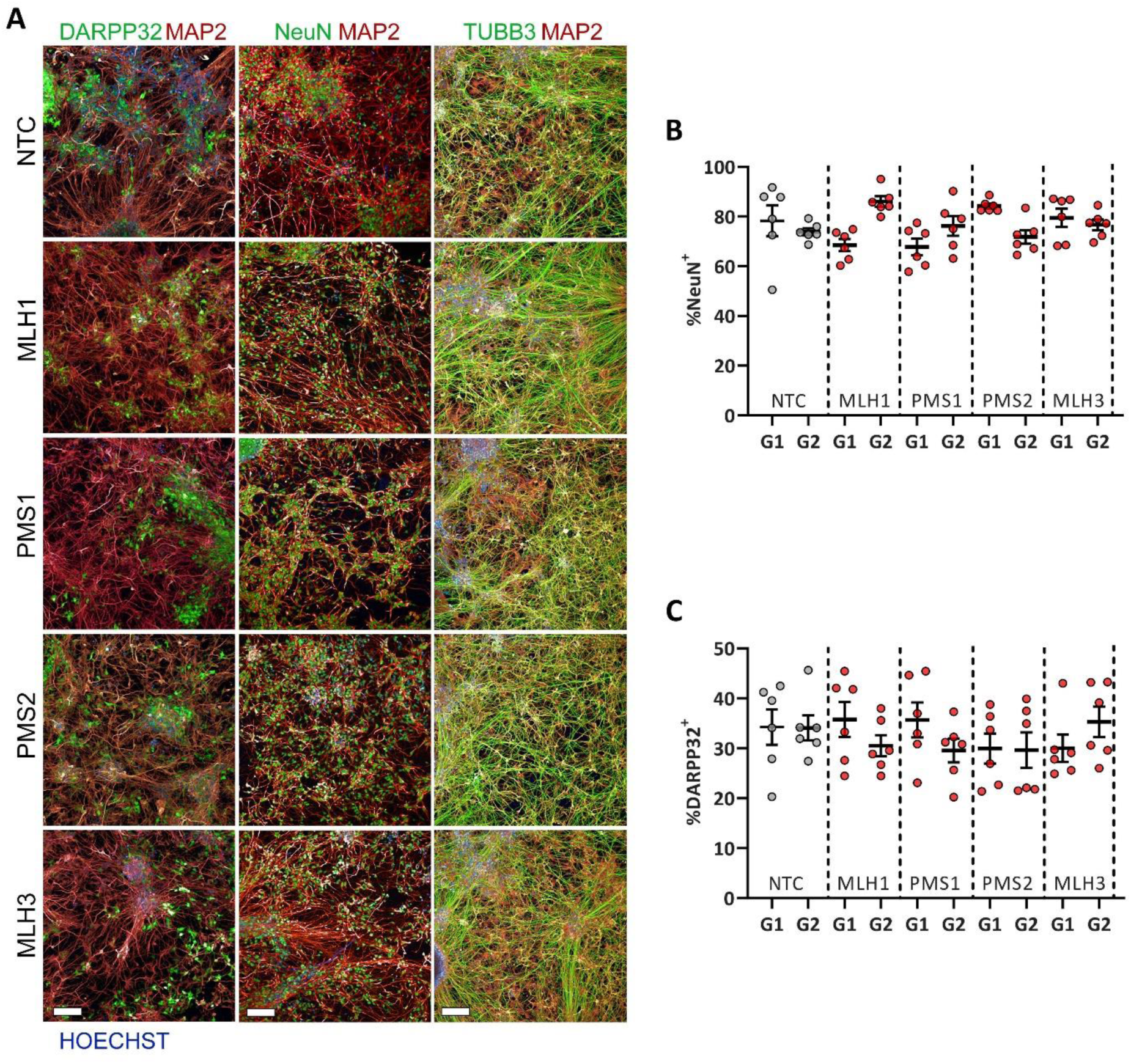
Constitutively lowering MMR factors does not affect iPSC competency to differentiate to a striatal identity. Representative immunostaining for one of each indicated guides for the neuronal markers NeuN, TUBB3 and MAP2, as well as the medium spiny neuron marker DARPP32. Scale bars 100µm. (A). Quantification of NeuN (B) and DARPP32 (C) positive cell frequency as proportion of the total cells counterstained with Hoechst. Five fields each from three replicate wells of two striatal differentiations with mean bar ±SEM for each targeting and non-targeting guide.

We used a timepoint thirty-six days post-initiation of differentiation as the baseline reference for assaying repeat expansion in striatal neurons over a nine-week period (Figure 6A). Transcript levels were assayed at week zero (baseline) and week nine and showed that target lowering was maintained over the course of the experiment relative to the non-targeting controls (Figure 6B).

**Figure 6.**
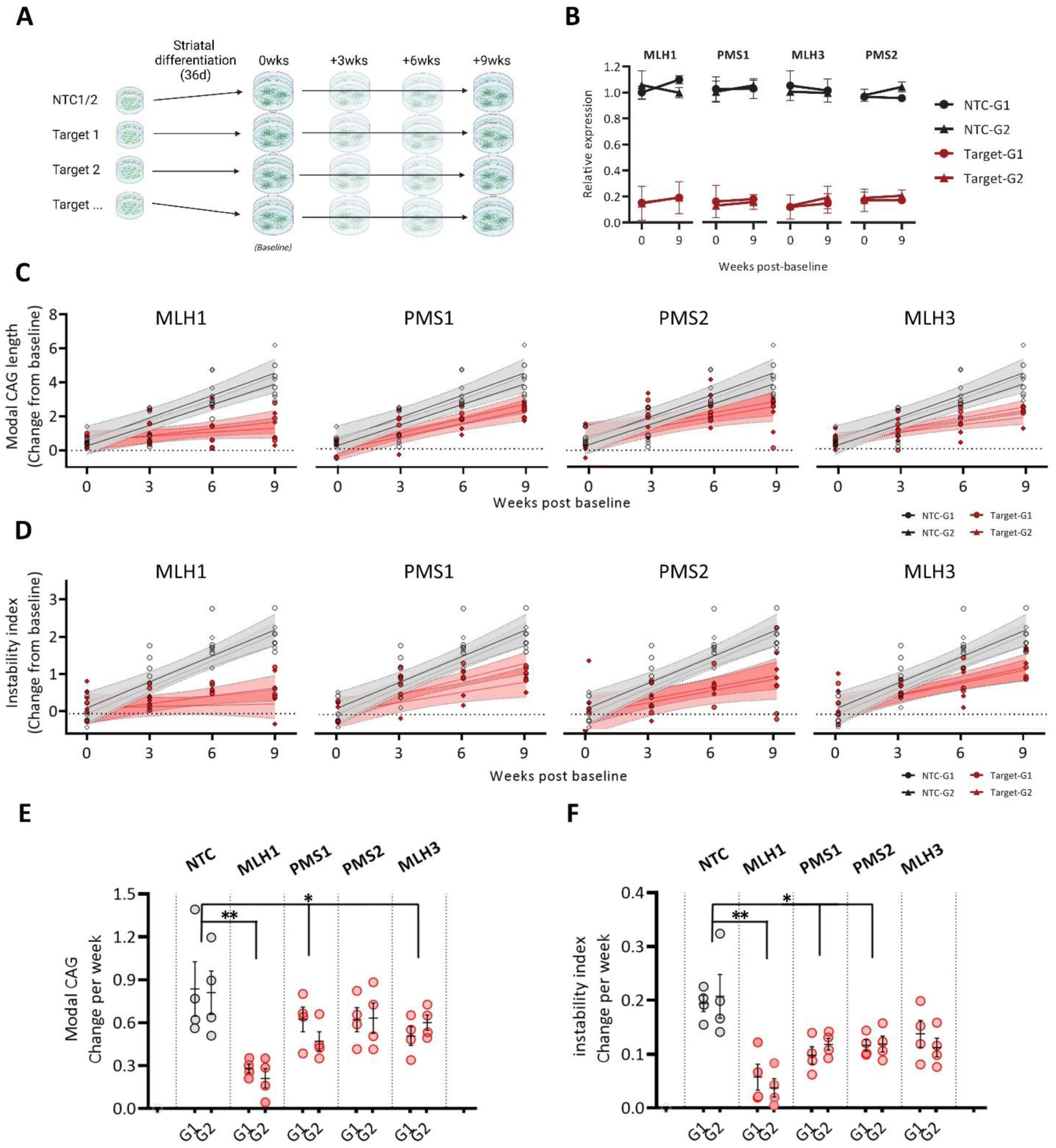
Reduced expression of each MutL factor slows somatic expansion in differentiated striatal neuron cultures. Somatic instability was measured in cultures of striatal neurons differentiated from MLH1, PMS1, PMS2, MLH3 and non-targeting guide CRISPRi iPSC pools. CAG length baseline is set at 36 days post neural induction (A). qPCR for each target shows lowering persists over the course. Mean expression relative to day zero ± SEM in triplicate for two NTC guides and the selected two target specific guides at baseline and week nine (B). Modal CAG repeat lengths over a nine-week period for MLH1, PMS1, PMS2 & MLH3 lowered pools in comparison to NTC relative to baseline (C). Change in instability index over a nine-week period for MLH1, PMS1, PMS2 & MLH3 lowered pools in comparison to NTC relative to baseline (D). Two differentiations of CRISPRi pools carrying one of two guides per target with four replicates. Dashed lines 95% confidence intervals. Rate of change in modal CAG per week (E) and change in instability index (F). Open circles represent parallel cultures with mean bar ±SEM. * P<0.05 * P<0.005.

The change from the baseline modal repeat length and instability index relative to controls were quantified over this nine-week period. The modal CAG length in control striatal cultures carrying the NTC guides increased by an average of 4-5 additional repeats (Figure 6C) and showed an increase in overall instability by almost 2 units (Figure 6D). Lowering MutL in striatal cultures led to a similar pattern of slowed repeat expansion as that seen in iPSCs. In cultures where MLH1 levels were lowered, around 1 repeat was added to the modal CAG length while reduced levels of PMS1, PMS2 and MLH3 reduced expansion to 2-3 repeats over the same time course (Figure 6C). Comparison of the rate of modal CAG changes shows reduced MLH1 expression slows repeat expansion by 69% (P<0.005), while targeting PMS1, PMS2 and MLH3 slow it by 25%, 21% and 28% respectively (P<0.05, Figure 6E).

These trends were maintained in the instability index (Figure 6D) – while lowering MLH1 had the largest impact on reducing instability, an effect was also observed where PMS1, PMS2 and MLH3 levels were reduced. In NTC striatal cultures, instability index increased by an average of 0.20 units per week and lowering MLH1 significantly reduced this by nearly 75% to around 0.05 units per week (P<0.005, Figure 6F). Lowering expression of either PMS1, PMS2 or MLH3 also resulted in a lower instability but to a lesser degree with reductions of 46%, 41% and 38% respectively compared to controls (P<0.05, Figure 6F).

### MMR factor levels are substantially reduced during differentiation to striatal neurons

While iPSCs maintain the necessarily robust DDR factor expression of hESC and embryonic inner cell mass (Lin et al., 2014), somatic cells typically express lower levels. This can be seen during the differentiation of the iPSCs to a striatal identity – loss of the pluripotency marker OCT4 and expression of both the post-mitotic neuronal marker NeuN and striatal marker CTIP2, is accompanied by substantial reductions in all MMR factor levels in differentiated cultures (Figure 7A).

**Figure 7.**
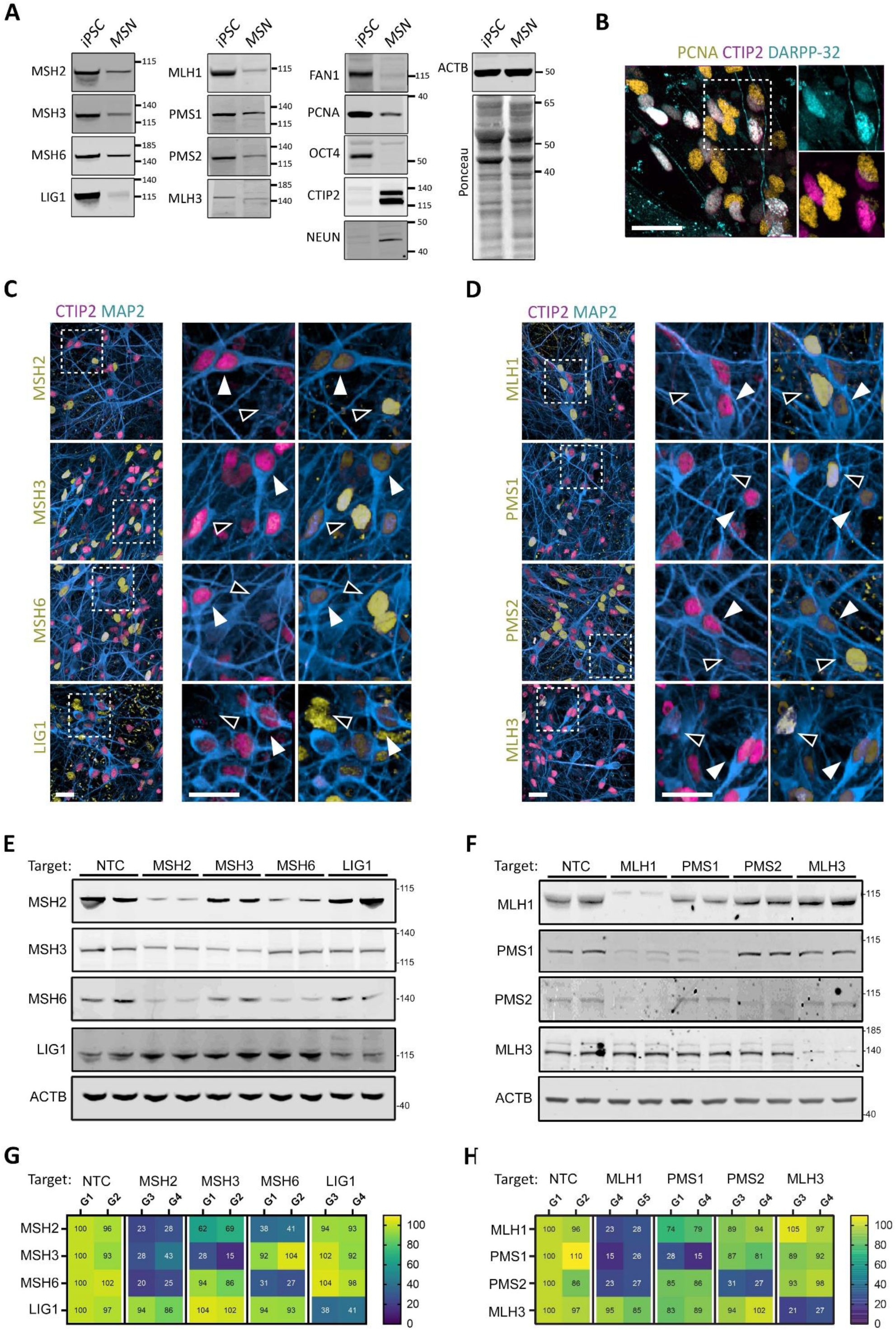
MMR factors are present at very low levels in medium spiny neurons and lowering expression impacts levels of partner proteins in iPSCs. Western blot comparison protein levels in 125Q CRISPRi HD iPSC before (iPSC) and after striatal differentiation at day 100 (MSN)(A). Immunostaining for PCNA, CTIP2 and DARPP-32 at intermediate stages of differentiation on day 40, dashed box indicates insets, scale 100µm (B). Immunostaining for CTIP2 and MAP2 in conjunction with the indicated MMR factor on day 100, dashed box indicates inset location, scale 100µm White arrow heads – CTIP2/MAP2^+^ MSNs, black arrow heads – MMR factor^+^ non-neuronal cells(C). Western blot for MMR proteins in undifferentiated iPSCs from CRISPRi pools lowering expression of either MutS & LIG1 (E), or MutL (F) from two separate guides (pairs of columns). Heat maps for the corresponding blots quantifying the effects of lowering expression of the indicated target on protein level. Mean of three blots, normalised to ACTB, and presented relative percentage of levels in the non-targeting control guide NTC-G1.

PCNA has long been implicated in MMR in dividing cells (Umar et al., 1996), and recent work by Phadte et al., 2023 has invoked PCNA as a requirement for CAG loop-out removal by FAN1 in competition with MutSβ using *in vitro* systems. In contrast to dividing cells, PCNA is not typically observed in healthy post-mitotic neurons (Ino and Chiba, 2000). While PCNA was observable in bulk by western blot (Figure 7A), this signal likely comes predominantly from the small proportion of non-neuronal cells also present. We confirmed by immunostaining that very low to undetectable levels of PCNA were present in CTIP2^+^/DARPP-32^+^ neurons while substantially higher expression was seen in non-neuronal cells at early time points during neurogenesis (Figure 7B, D40).

Similarly, expression of MutS and LIG1 (Figure 7C), and MutL (Figure 7D) are significantly lower in striatal neurons (CTIP2/MAP2^+^, white arrows) than non-neuronal cells (black arrows) in D100 cultures. LIG1 expression levels in striatal neurons appear higher but levels still remain less intense than that seen in non-neuronal levels in the same cultures (Figure 7C) Unlike the robust nuclear distribution seen for the other targets, MLH3 shows a distinct nucleocytoplasmic distribution as previously reported in somatic and cancer cell lines (Korhonen et al., 2007; Ou et al., 2009).

### MMR co-factor protein levels are altered in addition to targeted factors

The levels of each protein in many multiprotein complexes such as MutS and MutL, are regulated through the relative abundance of their component factors, with ‘orphan’ excess co-factors targeted for degradation (Juszkiewicz and Hegde, 2018). Therefore, to understand how the lowering of each target may influence repeat expansion we must also look at the remaining levels of the relevant co-factors in the MutS and MutL complexes (Abildgaard et al., 2019; Arlow et al., 2021; Cannavo et al., 2007). To characterise the extent to which altered co-factor abundance occurs because of targeted lowering, we quantified the protein levels of each MMR factor within each complex in iPSC cultures (Figure 7EF). These data were collected from iPSC cultures due to the heterogeneous levels of expression observed in the striatal neuron cultures.

Targeting MSH2 expression reduced MSH2 protein levels to ∼25% of control levels with either guide used (Figure 7EG). This reduction in MSH2 was accompanied by a substantial decrease in MSH6 levels as well as a smaller decrease in MSH3 levels. In contrast, where MSH3 lowering reduced MSH3 protein levels down to 15-29% of controls, the remaining MSH2 levels were reduced to only 62-69% and MSH6 levels remained unchanged. Targeted lowering of MSH6 resulted in a reduction of MSH6 protein to 31-27% of control levels, a larger loss of MSH2 was seen compared to that associated with MSH3 lowering, down to 38-41%. MSH6 lowering had no effect on MSH3 levels. The same pattern can be seen in U2OS knock-out cell lines. The loss of MSH6 resulted in a significant reduction in the level of MSH2 protein but left MSH3 unaffected in U2OS (Figure S2A). Similarly, in MSH3 null U2OS, MSH2 levels showed a limited reduction (Goold et al., 2021). Immunoprecipitation of MSH3 from MSH6 null U2OS showed the remaining levels of MutSβ were unchanged (Figure S5D). LIG1 levels in iPSCs were reduced to around 40% where directly targeted but remained unaffected by the lowering of any other target and did not affect their abundance either (Figure 7G).

A similar pattern of reduction in co-factor levels occurs where components of MutL are targeted in the HD iPSCs. Lowering MLH1 reduced protein level to 23-28% between the two guides used relative to control guides (Figure 7FH). This was accompanied by significant reductions in the levels of both PMS1 (15-26% remaining), PMS2 (23-27% remaining) but no significant effect was seen on MLH3 levels (Figure 7H). Again, this was also observed in U2OS MLH1 null cells where PMS2 is almost undetectable and MLH3 levels remain unaffected (Figure S2A). Lowering either PMS1 or PMS2 in iPSCs also resulted in a reduction of MLH1 levels, with a larger effect on MLH1 levels seen where PMS1 was lowered (Figure 7H).

## DISCUSSION

Genetic studies to identify disease modifiers in human populations also offer valuable insight into the therapeutically exploitable tolerability of reduced gene function. Using disease-relevant cell types we have shown MMR-associated modifiers of HD onset and progression play a role in repeat expansion dynamics at the expanded *HTT* CAG repeat, as do MMR-associated factors not highlighted in GWAS. We also show that lowering rather than complete ablation of gene expression is sufficient to attenuate repeat expansion in these models.

MutSβ has been shown to be an almost essential driver of somatic expansion in multiple models across multiple repeat expansion associated disorders including HD, and MSH3 locus modifiers are associated with delayed onset and progression (Flower et al., 2019; Moss et al., 2017). We confirm this also occurs in our human *ex vivo* models where lowering either MSH2 or MSH3 has a dramatic effect on repeat expansion in our HD iPSCs.

In contrast MSH6 (and so MutSα) has not been shown to affect somatic expansion in HD mouse models, though it has been implicated in preventing intergenerational contractions in CAG repeat length (Dragileva et al., 2009). In a mouse models of Myotonic Dystrophy Type 1 (DM1), loss of MSH6 led to increased expansion in non-neuronal tissue (van den Broek et al., 2002). Here we found no significant change in repeat expansion upon lowering MSH6. Although we were able to detect a significant MMR defect, we cannot discount the possibility MSH6 levels simply weren’t lowered sufficiently for a change in repeat expansion rate to become detectable. The same may be the case for LIG1 which also showed no significant effect upon lowering and functional redundancy may occur for LIG1 through other ligases (Gazy et al., 2019; Masani et al., 2016). Data from Lig1 mouse models suggest this redundancy is more effective where protein levels are reduced in comparison to deleterious LIG1 variants as Lig1 null mice fare better than Lig1 mutant mice (Bentley et al., 2002). This is consistent with one onset-delaying modifier haplotype in LIG1 resulting in a deleterious missense mutation while another is associated with increased expression (Lee et al., 2022).

The missense MLH1 mutation I219V identified by GWAS is classed as benign and no evidence for changes in transcript levels was found (Lee et al., 2019), nor does it appear to affect protein stability (Abildgaard et al., 2019). Despite this, the effect we observe upon lowering MLH1 expression is concordant with the reduced repeat expansion reported in multiple Mlh1 null mouse models of HD (Loupe et al., 2020; Pinto et al., 2013), and our previous work in a U2OS cell system (Goold et al., 2021). The same effect has also been seen on expansion of the GAA repeat in FRDA models (Ezzatizadeh et al., 2014; Halabi et al., 2018). One could ascribe this effect entirely to depletion of the total MutL pool in the absence of MLH1, however we found that independently lowering each MutL co-factor, PMS1, PMS2 and MLH3, also slowed repeat expansion to a lesser degree despite their reported functional differences.

Individually each of the MutL complexes have been shown by Miller et al., (2020) to be required for expansion of the Fmr1 CGG-tract in a mouse embryonic stem cell (mESC) model of fragile X-related disorders (FXD), however the consistency of effect between different cell and repeat types is not well characterised. A role for PMS1 in repeat expansion has not been previously examined either at the HD CAG repeat tract or in human models. Onset hastening and delaying modifier haplotypes were identified, though their effect on PMS1 function are unclear (Lee et al., 2019, 2022). Variants that are predicted to be deleterious to PMS1 function have been associated with a later age at onset and reduced disease severity in HD cohorts through exome sequencing (McAllister et al., 2022). Here we show for the first time lowering PMS1 levels slows CAG repeat expansion in human HD iPSC derived striatal neurons.

In keeping with each MutL complex being required for repeat expansion, we found that lowering MLH3 expression was also sufficient to slow expansion despite MLH3 abundance (and so MutLγ) being substantially lower than both MutLα and MutLβ here and in other somatic cell types (Cannavo et al., 2005). In FRDA & HD patient fibroblasts, a HD mouse model and an FRDA model cell line, endonuclease-dead MLH3 variants result in the abrogation of the relevant repeat expansion to the same degree as a complete knockout (Halabi et al., 2018; Hayward et al., 2020; Roy et al., 2021). Despite its clear role in repeat expansion, no modifier haplotypes are reported to associate with MLH3 in the GeM-HD GWASs (Lee et al., 2019). MLH3 variants are however associated with somatic instability in blood in HD populations (Ciosi et al., 2019). The lack of significance in the GWASs could be due to a low tolerance for reduced function MLH3 variants in germ cells during meiosis (Charbonneau et al., 2009; Dai et al., 2021).

We also show here that lowering PMS2 slows expansion of the *HTT* CAG repeat tract in both dividing iPSCs and post-mitotic striatal neurons. Again, modifier haplotypes have been associated both with early or delayed onset (Lee et al., 2019) and previous characterisations of PMS2 have reported roles in both suppression and promotion of expansion. These observations have been made primarily for the mouse ortholog Pms2 in the context of non-HD trinucleotide repeat tracts. Gomes-Pereira et al., (2004) found that loss of Pms2 slowed expansion of the CTG repeat in the brain of a DM1 mouse model and was accompanied by rare large contractions. Here we found lowering PMS2 slowed expansion by around 21% in line with Gomes-Pereira et al., (2004) who showed a 50% reduction in Pms2 null mice. A more dramatic effect was seen on the *Fmr1* CGG repeat where loss of Pms2 completely halted expansion in FXD mESCs (Miller et al., 2020). Conversely, Halabi et al., (2018) found no significant change in expansion rate of a construct carrying the FRDA GAA repeat in HEK cells where PMS2 levels were lowered by shRNA. Bourn et al., (2012) found that Pms2 supresses expansion in the brain of an FRDA mouse model – loss of Pms2 resulted in increased frequency of large expansions.

This apparent plasticity of PMS2’s role in somatic instability could be due to cell-type, repeat structure or MutL abundance. One potential explanation is the competition model suggested by Zhao et al., (2021) whereby changes in repeat substrate length and MutLα/β/γ stoichiometry result in favouring either expansion, contraction, or maintenance. Within this model the loss of PMS2 and so MutLα supresses expansion at increased repeat lengths, and the reduced levels here in our system may be sufficient to reach this theoretical threshold.

The implications of each MutL complex influencing repeat expansion rates is interesting mechanistically. PMS2 and MLH3, complexed with MLH1 as part of MutLα and MutLγ, are both independently capable of mediating repair in cell extracts otherwise lacking endogenous MutL (Kadyrova et al., 2020; Pluciennik et al., 2013). While this appears to suggest redundancy between them, PMS1 is unable to mediate repair in MutL deficient extracts. Loss of either Pms2 or Mlh3 in mouse leads to high tumour burdens despite Mlh3 being substantially less abundant than either Pms1 or Pms2 (Cannavo et al., 2005; Chen et al., 2005). Altering the relative levels of the different MutL complexes could affect the stoichiometry of MutS:MutL and MutLα:β:γ binding to DNA lesions (Bradford et al., 2020; Elez et al., 2012).

The role each of the different MMR complexes play at repeat structures is not well understood. Recent work by Zou et al., (2021) characterised the effects of each on genome instability in a large panel of isogenic knock-out hPSCs using whole genome sequencing. In their system, loss of MSH2, MSH6, MLH1, and PMS1 to a lesser degree, results in very similar insertion/deletion (indel) signatures with deletions frequently found at long repetitive sequences. In contrast, PMS2 favoured both insertions and deletions at the same class of sequence. How this data correlates with effect at locus specific expansions was not examined but these repair signatures highlight the distinct effect of losing PMS2 in comparison to the other MMR factors.

Interpreting the loss of any individual MutS or MutL target is mechanistically difficult due to differential secondary effects on the levels of their heterodimer partners. Where MSH2 levels are reduced by CRISPRi we observed a loss of a larger proportion of MSH6 than MSH3, as previously described in yeast and human cells (Arlow et al., 2021; Sakellariou et al., 2022). While MSH3 has been reported to function outside of MMR in other DDR pathways it is always as part of MutSβ (Eichmiller et al., 2018; Oh et al., 2023). This may suggest MSH3 has a higher affinity than MSH6 for MSH2, resulting in a bias towards MutSβ formation where MSH2 is lowered. Regardless, the net reduction in MutSβ here is still sufficient to attenuate expansion rates, as seen in heterozygous *Msh3* knock-out or *Msh3* siRNA treated mice (Dragileva et al., 2009; O’Reilly et al., 2023). The reciprocal effect was seen upon lowering either MSH3 or MSH6, however MSH2 levels were more substantially reduced where MSH6 levels were targeted, fitting with the higher abundance of MutSα than MutSβ (Reyes et al., 2015).

Similarly, where MLH1 levels are lowered it is intuitive to assume proportional lowering of the function of MLH1-containing MutL complexes. This is supported by the MMR deficiency observed in response to 6-TG. However, it is complicated by MLH3 stability being independent of MLH1 abundance (Here, and Cannavo et al., 2005), particularly in neurons as MutLγ (MLH1:MLH3) does not absolutely require PCNA for loading onto DNA or activity in cell-free systems (Kadyrova et al., 2020). This draws parallels to the stability of MSH3 upon lowering MSH2. However unlike MSH3, MLH3 (and PMS2) have been reported to function independently of the MLH1 complexes (Pannafino and Alani, 2021). These effects could be dissected by lowering multiple targets at once. As lowering each non-MLH1 MutL component slowed expansion to some degree it also raises the question as to whether these effects are synergistic and by targeting multiple components would we see additive expansion rate slowing.

Mätlik et al., (2023) have performed single cell transcriptomic and repeat sizing in post-mortem striatal tissue from HD and SCA3 donors, and have clearly shown MSNs are exquisitely vulnerable to CAG repeat expansion in both the mutant *HTT* and *ATXN3* allele. They found an MSN-specific increase in MSH2 and MSH3 protein and transcript levels. Here we found expression of MMR factors to be at lower levels in differentiated striatal neurons, in comparison to levels in undifferentiated iPSCs and non-neuronal cells present within the differentiated cultures. Seriola et al., (2011) found the same in hESC models of HD and DM1 upon differentiation to osteoprogenitor-like cells. Reduced MMR gene expression was accompanied by reduced repeat instability in the differentiated cells compared to their parent hESC. One corollary of hPSC derived models is the juvenile nature of neurons generated. Transcriptomics have shown iPSC derived neurons to be comparable to mid-gestational identities *in vivo* (Handel et al., 2016; Mehta et al., 2018). The high metabolic rate of neurons and associated increase of genotoxic byproducts leaves neurons particularly vulnerable to DNA damage (Hegde et al., 2012). DDR gene expression is increased in response to damage (Belloni et al., 1999; Tennen et al., 2013), and this increases alongside the accumulation of DNA damage in aging brains (Lombard et al., 2005; Lu et al., 2004). Oxidative stress has been linked to repeat expansion in R6/1 mESCs with impaired DSB repair (Jonson et al., 2013). Perhaps transient increases in MMR factor expression in response to DNA damage act to briefly licence expansion, a window which widens with aging.

Regardless of the mechanism, MSH3, PMS1, PMS2 and LIG1 are all tolerant of loss of function over a predicted two fold range and their variants are not readily removed from the population by selective pressures (Karczewski et al., 2017; Kosmicki et al., 2017). MSH2 and MLH1 are unlikely to be good therapeutic targets in the CNS as their loss is associated with significantly increased risk of CNS cancers, as is the loss of PMS2 to a lesser degree (Caccese et al., 2020; Sijmons and Hofstra, 2016). In addition to this, targeting the apparently flexible influence of PMS2 on repeat expansion could prove to be a double-edged sword without further investigation. We further highlight PMS1 as a potential therapeutic target for slowing repeat expansion. In this report we have shown that lowering expression of MMR factors identified as HD onset modifiers to levels regularly achieved by current therapeutics was sufficient to slow the pathogenic expansion of the *HTT* CAG repeat tract. This further lends support to the search for an intervention which could potentially delay onset and progression in HD and other repeat expansion disorders which demonstrate somatic instability.

## ACKNOWLEDGMENTS

S.J.T. received funding from the CHDI Foundation. She is also partly supported by the UK Dementia Research Institute, which is funded by the UK Medical Research Council, Alzheimer’s Society, and Alzheimer’s Research UK. R.F & R.G were funded by the CHDI foundation.

The U2OS cell line was kindly gifted by Prof. John Rouse (University of Dundee, Scotland).

We would like to thank the CHDI Foundation and UK Dementia Research Institute, which receives its funding from the UK Medical Research Council, Alzheimer’s Society, and Alzheimer’s Research UK. The authors thank all S.J.T laboratory members, as well as Thomas Vogt, Brinda Prasad, Michael Finley and other members of the CHDI foundation for helpful discussions.

## AUTHOR CONTRIBUTIONS

S.J.T. supervised and led the work. S.J.T. and M.F. conceived the project. R.F. designed the experiments and performed iPSC and neuronal culture and associated assays and analyses. R.G. performed all U2OS cell culture and associated assays and analyses. M.F and L.C. performed and analysed fragment analysis. S.J.T. and M.F secured funding. The manuscript was written by R.F., with editing by all co-authors.

