## Supplemental Material for "Therapeutic validation of MMR-associated genetic modifiers in a human *ex vivo* model of Huntington’s disease"

### SUPPLEMENTAL MATERIALS

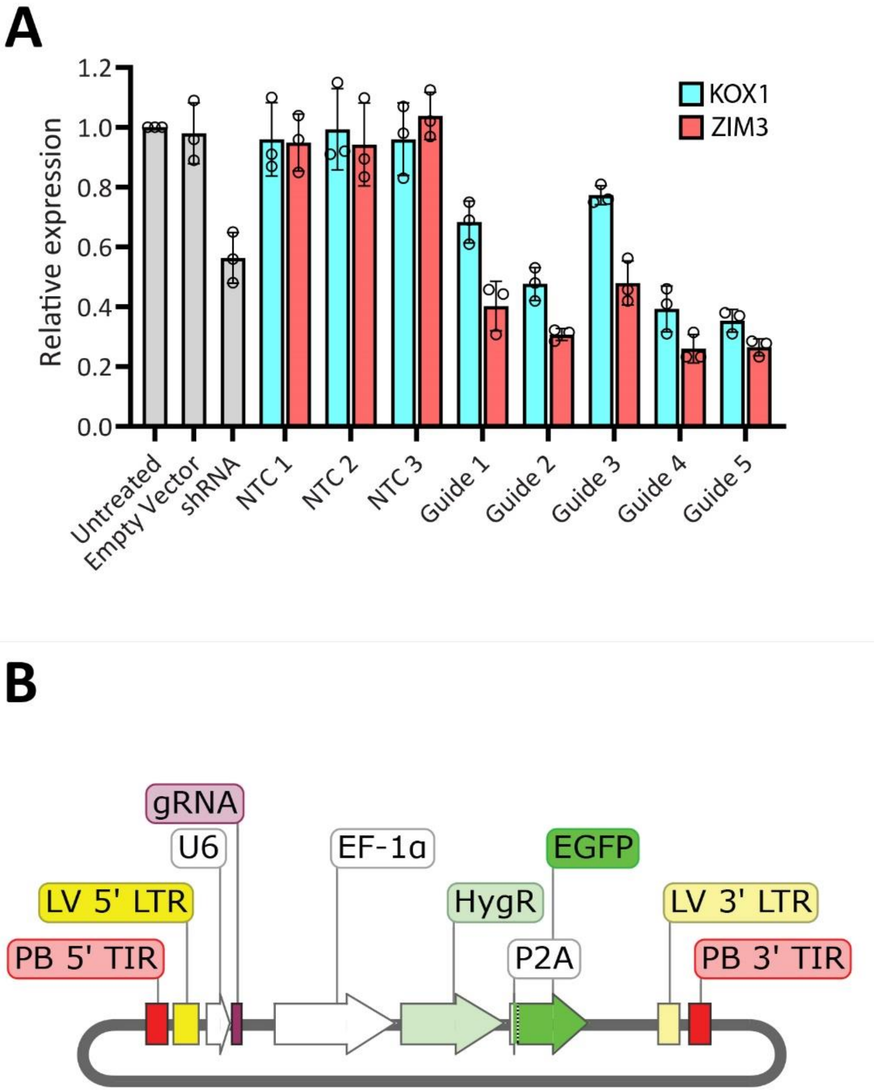


**Figure S1 – A more effective repressor construct and guide delivery vector.** Comparison between KOX1 and ZIM3 KRAB domains indicated better or same knock-down using PMS2 targeting guides (A). Guide sequences were cloned into either pLentiguide GFP-HYG or a modified version with additional PiggyBAC terminal inverted repeats (B).

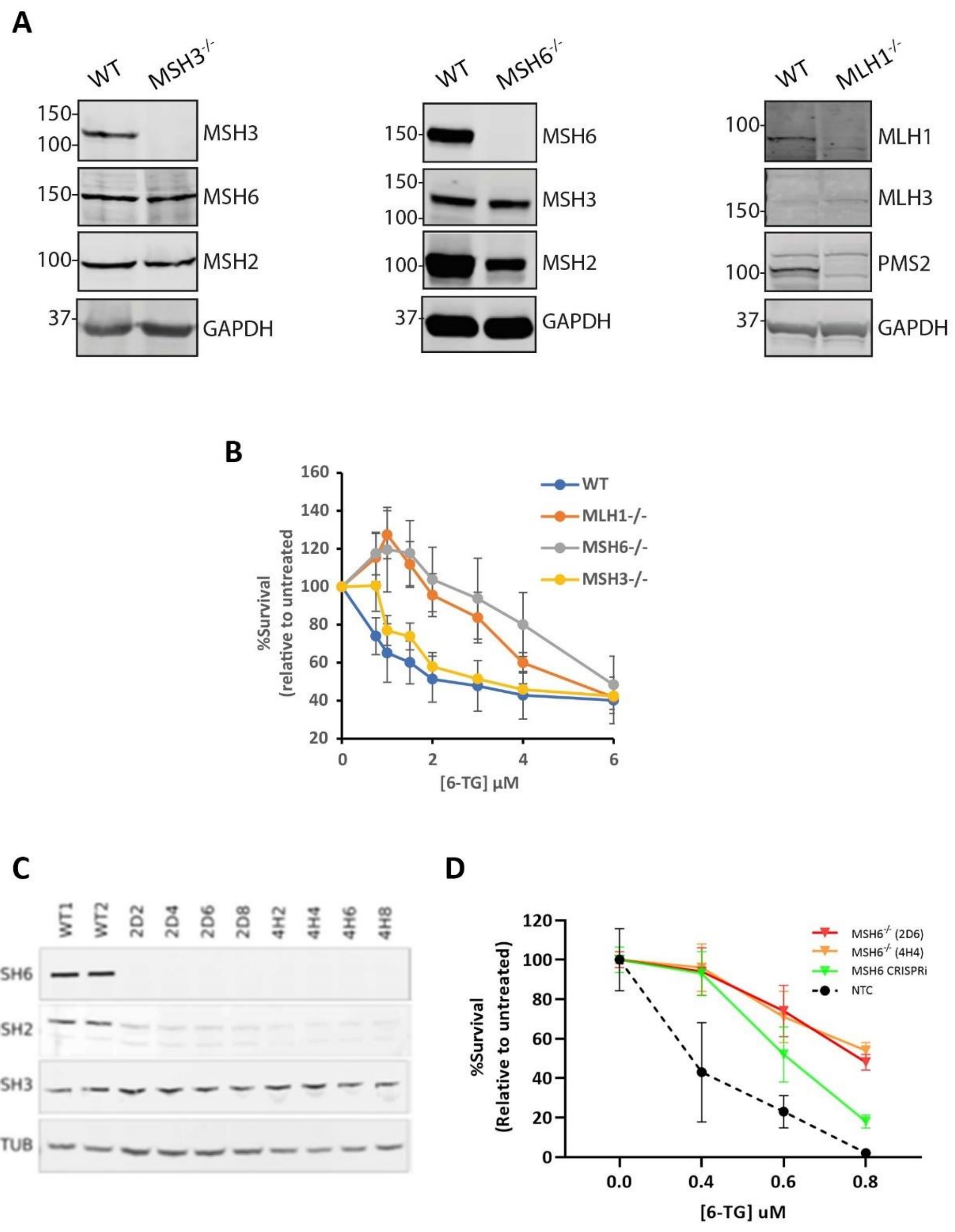


**Figure S2 – Knockout of MMR factors in U2OS cells results in MMR deficiency.**

Western blot for MMR associated proteins in U2OS MSH3, MSH6 & MLH1 knockout clones (-/-), compared to wild type U2OS (WT) (A). MMR deficiency seen after 6-thioguanine treatment of MSH3, MSH6 and MLH1 null and wildtype U2OS cells (B). Western blot of MSH6 knockout 125Q iPSCs generated using the same strategy with deletions in exon 2 or 4 (C). MSH6 knockout iPSC (clones 2D6 & 4H4) show further reduced sensitivity to 6-TG treatment than CRISPRi lowered MSH6 iPSCs (D).

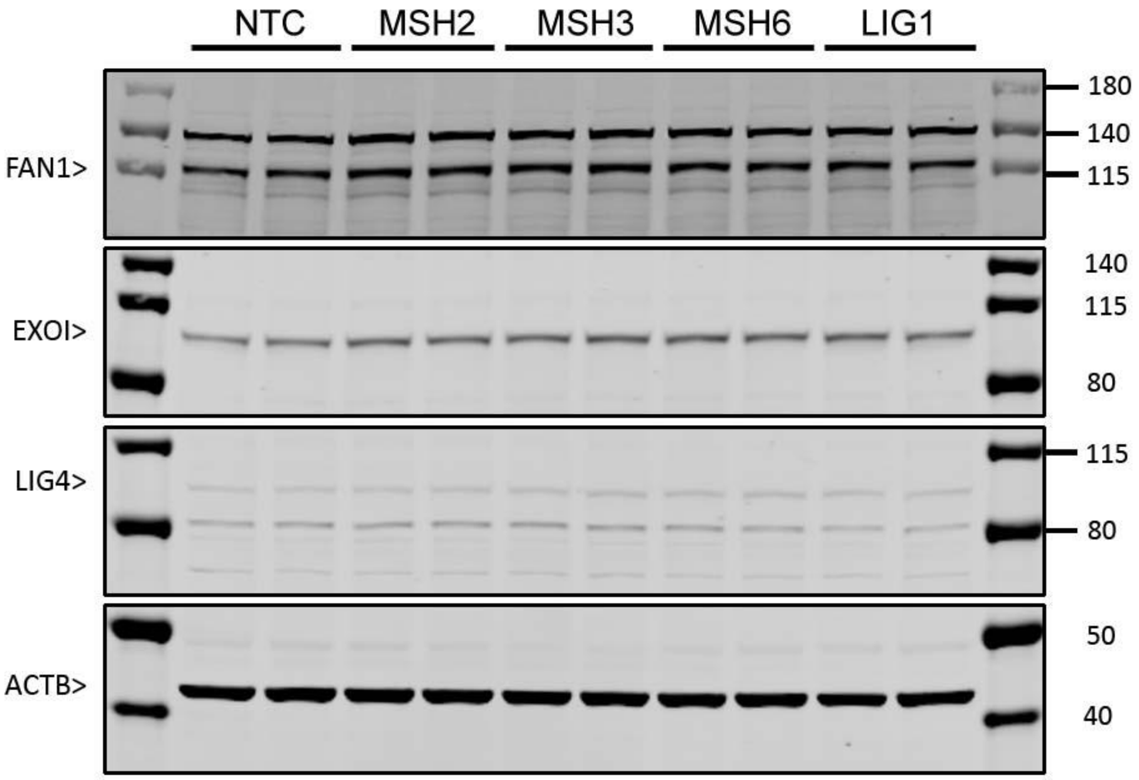


**Figure S3 – No changes seen in LIG4, EXO1 or FAN1 in CRISPRi MutS pools.** Western blots for indicated proteins (left) in two different CRISPRi pools targeting the indicated MutS factors and LIG1.

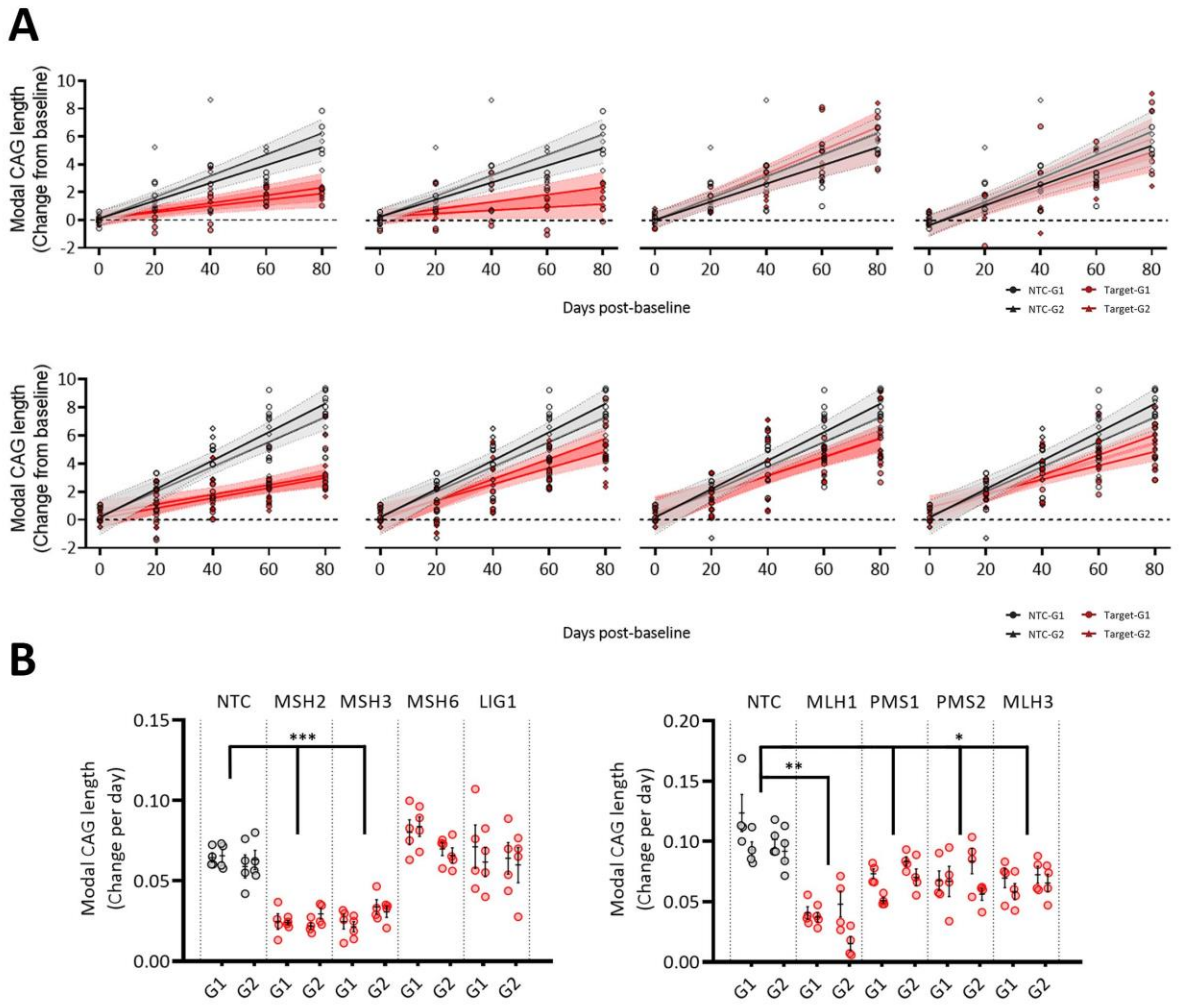


**Figure S4 - Reduced expression of MutS and MutL components slows increase in modal repeat length in dividing iPSCs.** Change in modal CAG repeat length over an eighty-day period for MutS & LIG1 (A), and MutL (B) lowered cultures relative to baseline on D0. Four cultures per guide passaged in parallel each from two independent CRISPRi pools, dashed lines 95% confidence intervals. Rate of change in modal repeat length per day for MutS & LIG1 (C), and MutL (D). Open circles represent parallel cultures with mean bar ±SEM. * P<0.05 * P<0.005.

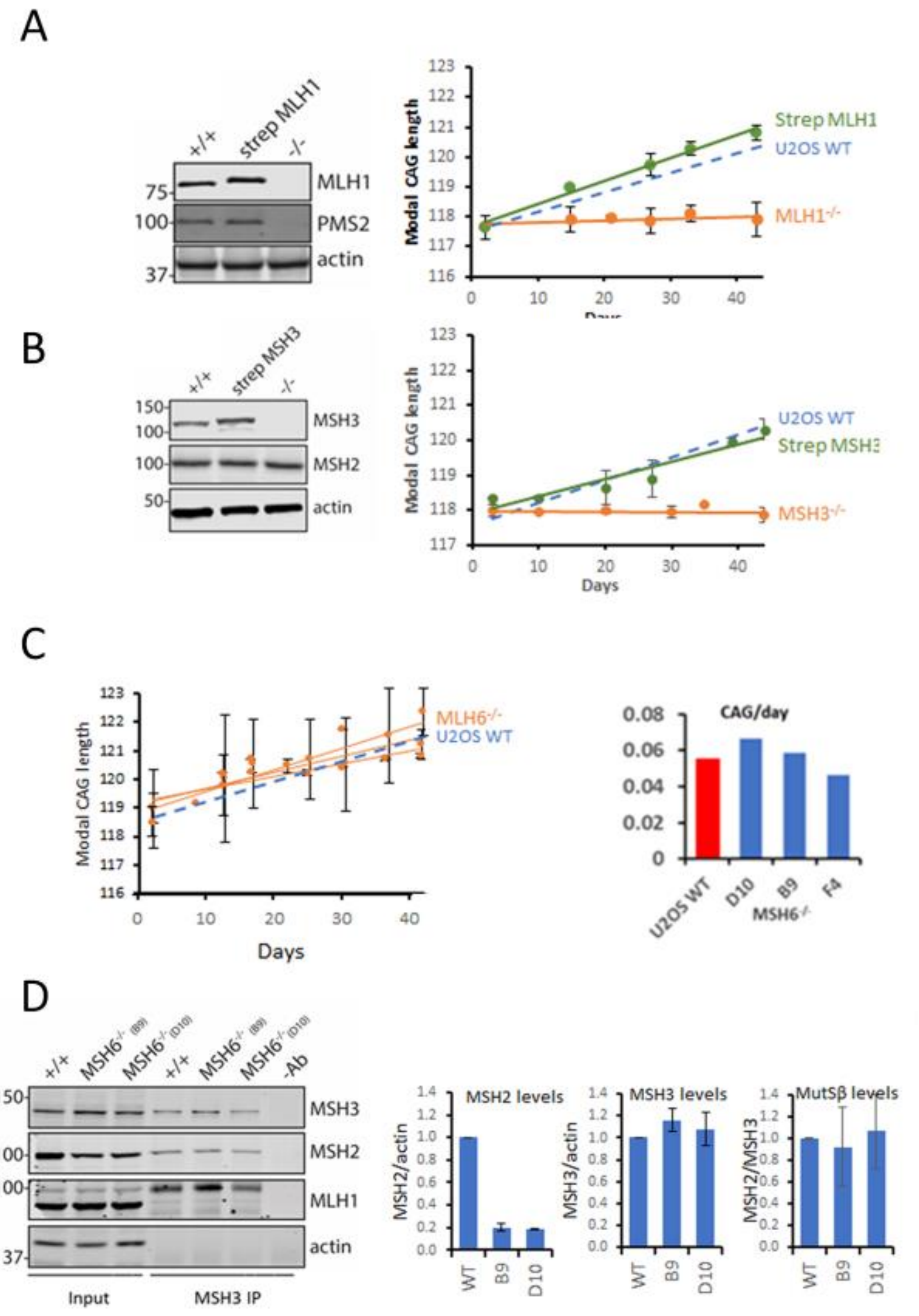


**Figure S5 – Repeat expansion in MSH3, MLH1 and MSH6 null U2OS.** A 118Q repeat construct introduced by transduction shows an increase in modal length over 40 days. Expansion is halted in MLH1$^{-/-}$ U2OS cells but can be rescued to WT U2OS cells rates by adding back a strep-tagged MLH1 to near physiological levels by dox-inducible expression (A). Expansion is halted and rescued in MSH3$^{-/-}$ U2OS cells using the same system (B). No change repeat expansion is observed in MSH6$^{-/-}$ U2OS cells (C). Immunoprecipitation shows that while MSH2 levels are lowered in MSH6$^{-/-}$ cells there is no change in MSH3/MSH2 interactions (D)

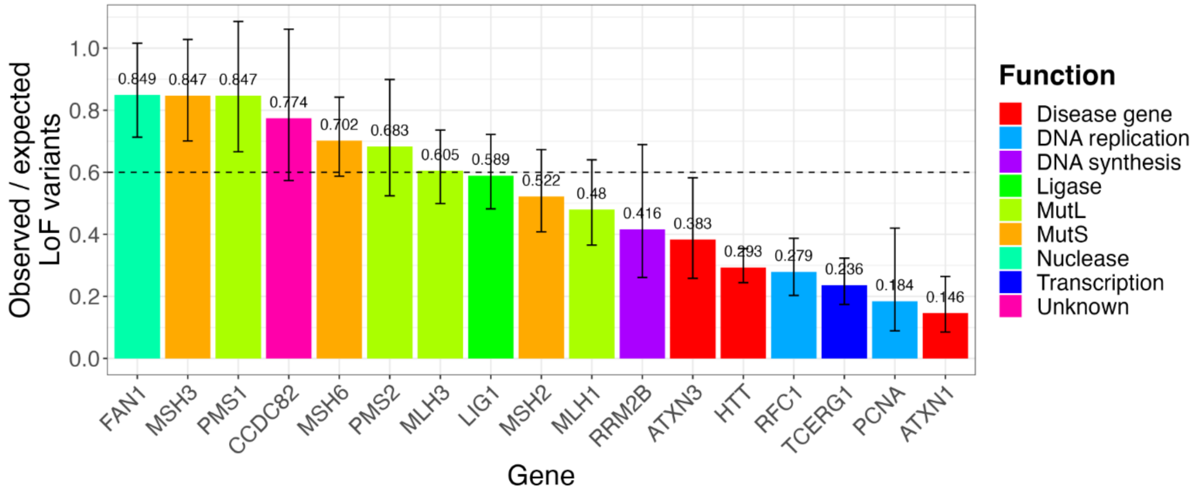


**Figure S6 – Genetic constraint in selected HD modifier and DNA repair genes.** Bar chart of loss-of-function metrics for selected genes, based on data from gnomAD (Gudmundsson et al., 2022). The y-axis represents the ratio of observed / expected ratio of loss-of-function variants (pLoF) for each gene, using GENCODE v39 annotations and GRCh38 reference. Error bars represent the 90% confidence interval. Lower o/e values indicate genes are under stronger selection. An o/e ratio of 0.2, for example, means there were 20% of the expected number of variants observed in that gene. Bars falling below the horizontal dashed line at an o/e ratio of 0.6 are generally considered to be under a high degree of genetic constraint, with significant selection against loss-of-function variation.

#### Table S1 – CRISPRi guide sequences

| Target | Guide | Sequence (5'-3') |
|---|---|---|
| **NTC** | 1 | TCTCTCGGAGTGGAGCAACA |
| **NTC** | 2 | GTCAGGTAATAGTCGGACTC |
| **NTC** | 3 | CGCAATCCCTTAGGATAGCC |
| **NTC** | 4 | ATCTACAATCCAGCCCTCTA |
| **MSH2** | 1 | GAGGTGAGGAGGTTTCGACA |
| **MSH2** | 2 | GTGAGGAGGTTTCGACATGG |
| **MSH2** | 3 | CGACATGGCGGTGCAGCCGA |
| **MSH2** | 4 | GACGCTGCAGTTGGAGAGCG |
| **MSH3** | 1 | AGGATGGCAGCCCGGCGGCA |
| **MSH3** | 2 | GGGCAAGGATGGCAGCCCGG |
| **MSH3** | 3 | GAGACATGGCAGGGCAAGGA |
| **MSH3** | 4 | GCTTCCGGCGAGACATGGCA |
| **MSH6** | 1 | AAAGCACCGCATCTACCGCG |
| **MSH6** | 2 | CAGGAGCCGCGCGGTAGATG |
| **MSH6** | 3 | GGCCCAACCGTTCTGTCGGA |
| **MSH6** | 4 | GCTCCGTCCGACAGAACGGT |
| **LIG1** | 1 | GCAGTCCCAAGTTCGCGCCA |
| **LIG1** | 2 | GAATGCCGTGGCGCGAACTT |
| **LIG1** | 3 | GCGCGAACTTGGGACTGCAG |
| **LIG1** | 4 | GCAACACACTCAGATCCGCC |
| **MLH1** | 1 | TCGGCGGCTGGACGAGACAG |
| **MLH1** | 2 | CCAAAATGTCGTTCGTGGCA |
| **MLH1** | 3 | CAAAATGTCGTTCGTGGCAG |
| **MLH1** | 4 | TCGTGGCAGGGGTTATTCGG |
| **MLH1** | 5 | TGGCGCCAAAATGTCGTTCG |
| **PMS1** | 1 | ACTACCTTCCTGCTAGCGCG |
| **PMS1** | 2 | GCGCTAGCAGGAAGGTAGTG |
| **PMS1** | 3 | ATTGCCTGCCTCGCGCTAGC |
| **PMS1** | 4 | GGCTGCTTGCGGCTAGTGGA |
| **PMS1** | 5 | CGCTGGCTGCTTGCGGCTAG |
| **PMS2** | 1 | CGGATCGGGTGTTGCATCCA |
| **PMS2** | 2 | AGCTGAGAGCTCGAGGTGAG |
| **PMS2** | 3 | CGAGCTCTCAGCTCGCTCCA |
| **PMS2** | 4 | ACGAGATCGCTGCAACACTG |
| **PMS2** | 5 | AACACTGAGGTCGCCACTCC |
| **MLH3** | 1 | AAGATCCAAGGTGCGCGCGT |
| **MLH3** | 2 | GTGTCGGAGAATTTGTTAAG |
| **MLH3** | 3 | TGTCGGAGAATTTGTTAAGC |
| **MLH3** | 4 | TTCTCCGACACCAACCGCCT |
| **MLH3** | 5 | TTCTCCGACACCAACCGCCT |

##### Table S2 – Primer sequences

| Primer | Sequence (5'-3') | Amplicon |
|---|---|---|
| **pLG_cPCR_F1** | TTTCAGACCCACCTCCCAAC | 2447bp +Stuffer |
| **pLG_cPCR_R1** | AGTGGATCTCTGCTGTCCCT | 558bp - Stuffer |
| **pLG_cPCR_F2** | ATCGTTTCAGACCCACCTCC | 2242bp +Stuffer |
| **pLG_cPCR_R2** | ATCGTTTCAGACCCACCTCC | 353bp - Stuffer |
| **PB 5P TIR - pLG** | cttcccaacagttgcCATGCGTCAATTTTACGCATGATTATCTTTAACGTACGTCACAATATGATTATCTTTCTAGGGttaaagcctgaatggcgaatgg | 1751bp |
| **PB 3P TIR - pLG** | ttcgccagttaatagtttgcCATGCGTCAATTTTACGCAGACTATCTTTCTAGGGTTAAgtctgacgctcagtggaacg | |
| **ZIM3_5P_HindIII** | cacaagcttaattctggctaactgtcggg | 398bp |
| **ZIM3_3P_Stop_MluI** | aattcacgcgtctaaaccactttgtacaagaaagttgggtagag | |
| **ZNF80_F1** | TGCAGCTCATCCTCACTTGG | 436bp |
| **ZNF80_R1** | GAGGCAAGGCCTTTGTACCT | |
| **GPR15_F1** | CTTGCATGAGTGTTGACCGC | 303bp |
| **GPR15_R1** | AATGGGCACACAGCTTCCTT | |
| **C13_5PJ_F1** | TATCGGTATCCTCGACTTGGC | 1843bp |
| **C13_5PJ_R1** | CTCCTCCACGTCACCGCA | |
| **C13_3PJ_F1** | GCAACCTGTTCAGTGCGTC | 1975bp |
| **C13_3PJ_R1** | AGTATGCTTATGCCAAGCCA | |
| **dCas9_F1** | CATCGAGCAGATCAGCGAGT | 275bp |
| **dCas9_R1** | CGATCCGTGTCTCGTACAGG | |
| **T7_MSH6_Ex2_F1** | GCCTTTTTCCTGCCATCAGC | 647bp |
| **T7_MSH6_Ex2_R1** | TTTCACAACTGCCACCCCTT | |
| **T7_MSH6_Ex4_F1** | TTGGCATATGAAGTTGCAGCATA | 642bp |
| **T7_MSH6_Ex4_R1** | GCTGTTCAGGCCTTCACTCT | |

#### Table S3 – Knockout guides

| MSH6 (NG_007111.1) | Sequence (5'-3') | Cut position |
|---|---|---|
| **Hs.Cas9.MSH6.1.AF** | AGCCUAAGACACAAGGAUCUGUUUUAGAGCUAUGCU | 20548 |
| **(Targeting)** | AGCCTAAGACACAAGGATCTAGG | |
| **Hs.Cas9.MSH6.1.AH** | AUUUAAGCCAGACACUAAGGGUUUUAGAGCUAUGCU | 20645 |
| **(Targeting)** | ATTTAAGCCAGACACTAAGGAGG | |

#### Table S4 – TaqMan Assays

| Target | Assay ID |
|---|---|
| **ATP5B** | Hs00969569_m1 |
| **EIF4A2** | Hs00756996_g1 |
| **UBC** | Hs00824723_m1 |
| **MLH1** | Hs00179866_m1 |
| **PMS1** | Hs00922262_m1 |
| **PMS2** | Hs00241053_m1 |
| **MLH3** | Hs00998142_m1 |
| **MSH2** | Hs00953527_m1 |
| **MSH3** | Hs00989003_m1 |
| **MSH6** | Hs00943000_m1 |
| **LIG1** | Hs01553527_m1 |

**Table S5 – Antibodies**

| Primary | Supplier | Cat# | Dilution |
|---|---|---|---|
| **MLH1** | BD | #554073 | 1:500 (ICC), 1:1000 (WB) |
| **PMS1** | Invitrogen | PA5-86724 | 1:500 (ICC), 1:1000 (WB) |
| **PMS2** | Invitrogen | PA5-87127 | 1:500 (ICC), 1:1000 (WB) |
| **MLH3** | Santa Cruz | sc-25313 | 1:500 (ICC), 1:1000 (WB) |
| **MLH3** | ProteinTech | 25298-1-AP | 1:500 (ICC), 1:1000 (WB) |
| **MSH2** | Cell Signalling Technologies | #2017 | 1:500 (ICC), 1:1000 (WB) |
| **MSH3** | ProteinTech | 22393-1-AP | 1:500 (ICC), 1:1000 (WB) |
| **MSH6** | BD | 610918 | 1:500 (ICC), 1:1000 (WB) |
| **LIG1** | ProteinTech | 18051-1-AP | 1:500 (ICC), 1:1000 (WB) |
| **FAN1** | University of Dundee / CHDI | FS2 | 1:2000 (WB) |
| **EXO1** | ProteinTech | 16253-1-AP | 1:2000 (WB) |
| **LIG4** | ProteinTech | 12695-1-AP | 1:2000 (WB) |
| **ACTB** | Abcam | ab8226 | 1:5000 (WB) |
| **TUB** | Abcam | ab6046 | 1:5000 (WB) |
| **GAPDH** | Abcam | ab9485 | 1:5000 (WB) |
| **CTIP2** | Abcam | ab18465 | 1:500 (ICC), 1:1000 (WB) |
| **DARPP-32** | Cell Signalling Technologies | #2306 | 1:500 (ICC) |
| **MAP2** | Novus | NB300-213 | 1:10,000 (ICC) |
| **NFL** | Cell Signalling Technologies | #2837 | 1:2000 (WB) |
| **TUBB3** | Abcam | ab41489 | 1:2000 (ICC) |
| **NEUN** | Abcam | ab104224 | 1:500 (ICC), 1:1000 (WB) |
| **PCNA** | Cell Signalling Technologies | #2586 | 1:500 (ICC), 1:2000 (WB) |
| **OCT4** | Santa Cruz | sc-5279 | 1:1000 (ICC), 1:1000 (WB) |
| **NANOG** | abcam | ab21624 | 1:1000 (ICC) |
| **SSEA4** | Invitrogen | MA1-021 | 1:1000 (ICC) |
| **LIN28** | Cell Signalling Technologies | #3978 | 1:1000 (ICC) |

| Secondary | Supplier | Cat# | Dilution |
|---|---|---|---|
| **IRDye 800CW Goat anti-Mouse (H+L)** | LiCor | 926-32210 | 1:15000 |
| **IRDye 680RD Goat anti-Rabbit (H+L)** | LiCor | 926-68071 | 1:15000 |
| **AlexaFluor 488 Gt α Ms** | Invitrogen | A-11001 | 1:2000 |
| **AlexaFluor 594 Gt α Rbt** | Invitrogen | A-11008 | 1:2000 |
| **AlexaFluor 488 Gt α Rbt** | Invitrogen | A-11012 | 1:2000 |
| **AlexaFluor 568 Gt α Rbt** | Invitrogen | A-11011 | 1:2000 |
| **AlexaFluor 488 Gt α Rt** | Invitrogen | A-11006 | 1:2000 |
| **AlexaFluor 647 Dk α Ms** | Invitrogen | A-31571 | 1:2000 |
| **Hoechst 33342** | Invitrogen | H3570 | 1:2000 |
